# Mechanosensitive dynamics of lysosomes along microtubules regulate leader cell emergence in collective cell migration

**DOI:** 10.1101/2022.08.03.502740

**Authors:** Rituraj Marwaha, Simran Rawal, Purnati Khuntia, Sanak Banerjee, Diya Manoj, Manish Jaiswal, Tamal Das

**Affiliations:** Tata Institute of Fundamental Research Hyderabad (TIFR-H), Hyderabad – 500 046, India

**Keywords:** Collective cell migration, Leader cells, Lysosomes, Actin dynamics, Epithelial tissue, Mechanotransduction

## Abstract

Collective cell migration during embryonic development, wound healing, and cancer metastasis entails the emergence of leader cells at the migration front. These cells with conspicuous lamellipodial structures provide directional guidance to the collective. Despite their physiological relevance, the mechanisms underlying the emergence of leader cells remain elusive. Here we report that in diverse model systems for wound healing, including cultured epithelial monolayer, *Drosophila* embryo, and mouse embryonic skin, leader cells display a peripheral accumulation of lysosomes. This accumulation appears essential for leader cell emergence, involves lysosomal movement along microtubules, and depends on the actomyosin contractility-generated cellular forces. Peripheral lysosomes associate with inactive Rac1 molecules to remove them from the leading periphery, which increases local Rac1-activity, triggering actin polymerization and promoting lamellipodium formation. Taken together, we demonstrate that beyond their catabolic role, lysosomes act as the intracellular platform that links mechanical and biochemical signals to control the emergence of leader cells.

## INTRODUCTION

Collective cell migration is the coordinated movement of cells, which is essential for organogenesis, embryo development, wound healing, tissue regeneration, and cancer metastasis^1–16^. For successful migration of epithelial cell collectives, a few cells at the migration front undergo phenotypic transformation to form distinct lamellipodial structures and emerge as leader cells^3, 17^. These cells provide guidance to collective cell migration^16, 18, 19^ and are critical for epithelial wound healing^2, 20^, branching morphogenesis^21, 22^, and cancer metastasis^1, 12, 23, 24^. Despite this physiological importance, what factors selectively designate only a few cells at the migration front as the leader cells remain largely unknown. Nevertheless, there are pieces of evidence that both mechanical and biochemical cues might be influential here^25, 26^. Mechanical cues such as the spontaneous fluctuations in the cellular force field, which is defined by the cell-cell and cell-matrix forces, influence the leader cell formation^25^. Much before the leaders start showing their characteristic lamellipodial structures, elevated traction forces and tensile stresses appear behind the would-be leader cells^25^. These results suggest a non-cell autonomous regulation of leader cell emergence. In addition to this mechanical regulation, spatial patterns of cellular signalling proteins such as p120-catenin^27^, Notch-Dll4^28^, and RhoA^26^ distinguish the leader cells from other non-leader cells at the migration front and the follower cells behind the leaders. The intrinsic polarized activity of Rac and RhoA GTPases is an integral signature of leader cell emergence^26^. Cues for leader cell emergence can also come from the extracellular milieu through the interaction of the migrating cells with the compliant extracellular matrix via focal adhesions^3^ and with the soluble signalling via self-generated chemokine gradient^29^. Furthermore, a recent study has identified the tumour suppressor protein p53 as a key marker for leader cells^2^. Elevated levels of p53 guide leader cell emergence and modulates the p21-CDK pathway to regulate collective migration^2^. But a connection between the mechanical cues and the biochemical events leading to leader cell emergence remains elusive.

As the sedentary cells become motile and develop a clear front-to-rear polarity, the intracellular organization undergoes a conspicuous transformation^3^. This transformation requires both breaking of the existing structures and the making of new structures. In this respect, previous studies have primarily focused on elucidating the dynamics of the actin cytoskeleton during the emergence of leader cells. However, the role of other intracellular structures, including many cellular organelles, in this process remains unexplored. Relevantly, one may presume that lysosomes, which are crucial for the catabolism and recycling of various biomolecules, might play a critical role in the cellular transformation to a motile state. Indeed, lysosomes can act as a signalling hub regulating cell growth, death, and differentiation^30^. They also house various proteins which regulate cell adhesion^31, 32^ and migration^33^. At a single-cell level, lysosomes regulate the dynamics of focal adhesions in migrating fibroblasts^32^. During dendritic cell migration, lysosomes accumulate in the direction opposite of migration and locally release calcium to modulate actomyosin dynamics supporting cell propulsion^34^. Finally, lysosomes respond to the mechanical properties of the extracellular matrix by altering their intracellular position^35^. While it seems that lysosomes can integrate mechanical signals and biochemical signals, very little is known about the specific roles of this organelle in collective cell migration. In this work, we explore the roles of lysosomes in the emergence of leader cells during collective cell migration.

## RESULTS

### Lysosomes accumulate at the leading periphery of the leader cells in various wound healing models

Previous research endeavours have investigated the biochemical, molecular, and cellular mechanisms underlying collective cell migration during the wound healing process, employing both *in vivo* and *ex vivo* model systems. For instance, stage-15 *Drosophila melanogaster* embryos have served as valuable *in vivo* models^36^, while mouse embryonic skin wounds have been utilized as *ex vivo* systems^37, 38^. It is known that the epidermal wound closure in stage-15 *Drosophila* embryos occurs by a combination of pursed-string contraction of actin cables and lamellipodial crawling^36^. First, to study the localization dynamics of lysosomes in wound healing, we used glass needles (0.5 µm inner diameter and 100 µm outer diameter) mounted on a femtojet microinjection system to create wounds in the stage-15 embryos. Since depending upon the size of the wound, the wound was closed completely in 1-2 hours, we fixed these embryos 15 minutes post-wounding to capture an intermediate stage of wound closure. We then stained these embryos for lysosomes and actin, using an antibody against a bonafide lysosome marker protein, LAMP1, and fluorescence-labelled phalloidin, respectively (Figs. 1A-B and Supplementary fig. 1A) and observed a few cells at the migration front emerging as leader cells having lamellipodial structures (Fig. 1B and Supplementary fig. 1A). Interestingly, the images revealed a distinct lysosomal accumulation at the cell periphery in these cells (Fig. 1B and Supplementary fig. 1A). To further verify the generality of our finding, we also analysed the localization of lysosomes in mouse embryonic skin wounds. We subjected the mouse embryonic skin samples dissected out of day 16.5 embryos to incisions and allowed the wound to heal for 30 mins in an air-media interface at 37℃, 5% CO_2_ humidified chamber (Fig. 1C). We then fixed and stained these samples for lysosomes and actin (Figs. 1C-D and Supplementary fig. 1B). Similar to the *Drosophila* embryos, here also we observed the movement of cells into the wound gap of the skin tissue and observe peripheral lysosome distribution in these peripheral migrating cells (Fig 1D and Supplementary fig. 1B). These results suggested that lysosome accumulation to the periphery of cells at the leading edge of wounds might be evolutionarily conserved.

**Figure 1:**
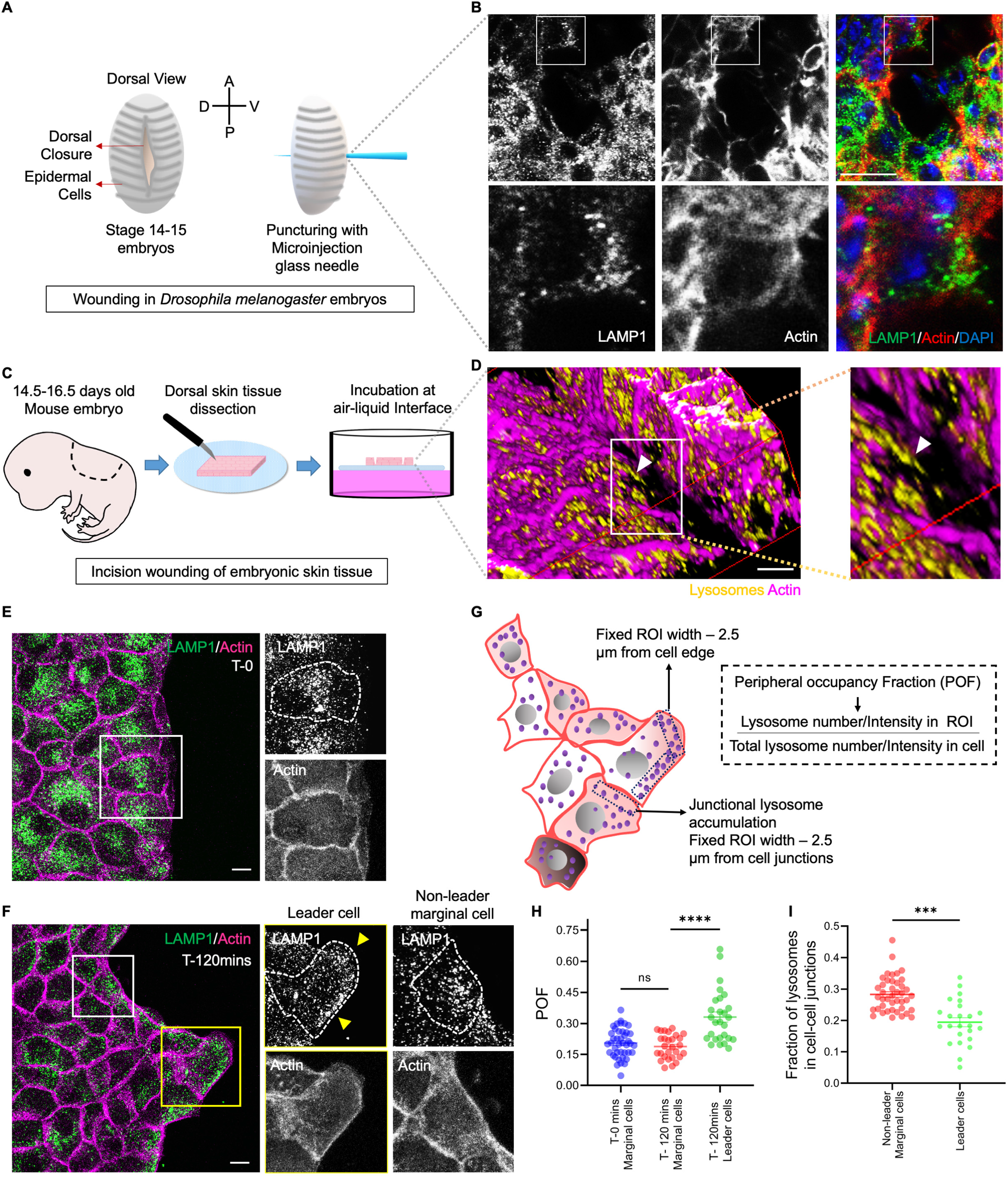
Enhanced lysosome accumulation to cell periphery in emerging leader cells during collective epithelial migration. **A)** Diagram representing stage-15 *Drosophila melanogaster* embryo wounding assay using a glass needle. **B)** Confocal micrographs of wound closure in *Drosophila melanogaster* embryos fixed 15 mins post-wounding and immunostained with Anti-LAMP1. Actin is labelled using Alexa-fluorophore conjugated phalloidin and DAPI stains the nucleus. The inset shows peripheral lysosome localization in cells migrating into the wound gap. **C)** Diagrammatic representation of skin epithelium dissection, wounding, and wound healing setup from day 16.5 mouse embryos. D) 3-Dimensional image representation of mouse embryonic skin wound post 30 mins of the incision. The wounded skin was PFA fixed and stained for lysosomes using Anti-LAMTOR-4 and actin was marked with Alexa-fluorophore conjugated phalloidin. **E-F**) Representative images of a migrating monolayer of MDCK cells fixed at T-0 (**E**) and T-120mins (**F**) post lifting of the confinement. Cells are immunostained for lysosomes using Anti-LAMP1 and Alexa-fluorophore conjugated phalloidin labels actin. Yellow arrowheads in (**F**) mark regions of lysosome accumulation in an emerging leader cell. **G**) Schematic representation of Peripheral Occupancy Fraction (POF) analysis, which measures peripheral lysosome accumulation. Junctional lysosomal fraction was measured by adding the lysosome number or intensity in all junctional ROIs and dividing by the total lysosome number/intensity. **H**) Scatter plot showing POF for non-leader marginal and leader cells at T-0 and T-120mins in MDCK cells (n=3, 15 cells analysed for marginal cells at T-0 and T-120mins; ∼10 leader cells per experiment were analysed). POF calculated from lysosome number **I**) Junctional occupancy fraction of lysosomes was calculated for non-leader marginal and leader cells, as represented by the scatter plot (n=3, 15 cells non-leader marginal cells and ∼10 leader cells analysed per experiment). Statistical significance was assessed using an unpaired Student’s *t*-test with Welch’s correction (two-tailed). Scale bars, 10 µm for all sub-figures; *** and **** denotes p-value<0.001 and p<0.0001, n.s is not significant.

Having discovered the peripheral accumulation of lysosomes in two model systems, we switched to a gap closure assay using cultured epithelial monolayer to perform live-cell imaging of the lysosome dynamics, investigate its actual role in the leader cell emergence, and elucidate the underlying molecular mechanism. To this end, we first grew Madin-Darby canine kidney (MDCK) epithelial cells to confluence into the wells of a migration chamber (Supplementary fig. 1C). In this system, after lifting off the confinement, cells migrated as a two-dimensional epithelial sheet. Leader cells emerged from the collectively migrating monolayer (Supplementary fig. 1D) at 2 hours post confinement lift-off. The leader cells, moving in front of the migrating monolayer, displayed extended lamellipodia, a crucial actin structure, which helped us mark these cells, and enlarged cell size^28^. To further investigate the dynamics of lysosomes in migrating epithelial monolayers, we fixed the migrating epithelial monolayer samples at 0, 15, 30, 60, 120, and 240 minutes post-confinement lift-off and immunostained them for lysosomes and actin (Supplementary fig. 2A). We observed that in the cells at the migration front (marginal cells), lysosomes accumulated at the cell periphery, whereas the cells in the follower layer did not show any change in lysosome localization (Supplementary fig. 2A) indicating that lysosome accumulation to the cell periphery in cells at the wound margin increases in a time-dependent manner. Interestingly, among the marginal cells, we observed further enhanced lysosome accumulation at the leading periphery of the leader cells, which was marked by their extended lamellipodial structures (Figs. 1E-F). We subsequently computed the fraction of peripherally localized lysosomes relative to the total lysosome number/intensity in a cell and reported it as Peripheral Occupancy Fraction (POF) (Fig. 1G). This quantitative analysis revealed a progressive increase in POF for lysosomes in cells at the margin as the migration progresses with time (Supplementary fig. 2B). A comparison of POF between the leader and non-leader marginal cells revealed a significantly higher fraction of lysosomes localized to the cell periphery in leader cells than in non-leader cells (Fig. 1H). Also, a closer examination of the distribution of lysosomes in leader and non-leader marginal cells revealed a lower junctional lysosome pool in leader cells as compared to non-leader marginal cells (Fig. 1I). To test the generality of our findings among cultured epithelial monolayer models, we next repeated the measurements in a murine mammary epithelial cell line, EPH4. Enhanced peripheral localization of lysosomes in leader cells emerged in this model as well (Supplementary figs. 2 D-E). Since leader cell emergence and lysosomal accumulation in these cells occurred at a similar time scale, we wondered how these dynamics behaved at the onset of collective migration in epithelial monolayers. Subsequently, we asked whether there might be any correlation between lysosome accumulation at the cell periphery in the marginal cells and collective cell movement. Intriguingly, the heat map of two events (Supplementary fig. 2E) showed a temporal correlation between collective cell movement and POF for lysosomes with different migration time points. As the epithelial cells migrated progressively, we observed an increase in lysosome accumulation starting at T- 30mins onwards and peaking at T-120mins post confinement lift-off.

Finally, to understand how the lysosome dynamics are connected to actin dynamics in leader cells, we performed live-cell imaging of LifeAct-eGFP-expressing MDCK cells having Dextran-647 (Alexa fluor 647-labelled dextran) pre-labelled lysosomes (Fig. 2A and Supplementary video 1). Lysosomes in these cells showed peripheral accumulation followed by dispersal, and subsequently, the lamellipodial protrusions started emerging. Distance versus time plots for leaders and other non-leader marginal cells showed differential lysosome movement along the cell periphery (Fig. 2B). Further, live-cell imaging of LifeAct-expressing MDCK cells having Dextran-647-labelled lysosomes clearly show that lamellipodia extension happened in lysosome-rich zones (Figs. 2C, D, and E-top panel and Supplementary video 2). In contrast, the actin structure was retracted from lysosome-deficient regions (Fig. 2E, bottom panel and Supplementary video 2). Taken together these results revealed that lysosome accumulation at the periphery of the leader cells is a ubiquitous event during collective cell migration across different model systems and is spatiotemporally correlated with the initiation and extension of lamellipodial structures in the leader cells.

**Figure 2:**
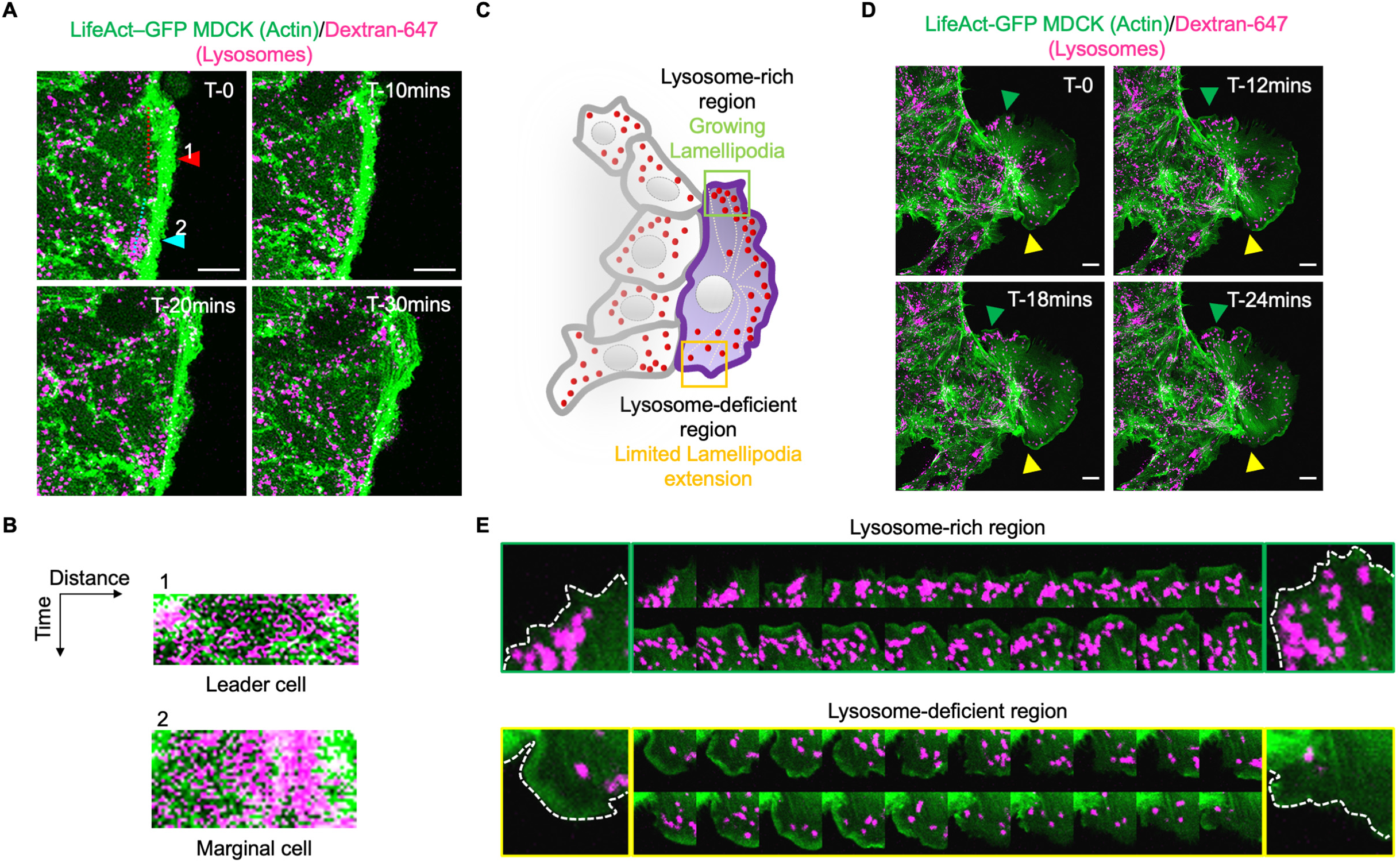
Lysosome accumulation to the cell periphery in leader cells promote lamellipodia formation. **A**) Live-imaging snapshots of actin and lysosome (labelled using Dextran-647) dynamics in LifeAct-GFP MDCK cells. The red arrowhead marks an emerging leader cell, and the cyan arrowhead marks a non-leader marginal cell. **B**) Kymographs showing differential lysosome dynamics in leader cell (1) and marginal cell (2) marked by red and cyan arrowheads respectively in A). **C**) Schematic representation of the relation between lysosome localization and lamellipodial dynamics in a leader cell during collective epithelial migration. **D**) Live imaging snapshots of lamellipodial and lysosome dynamics in a leader cell. White and yellow arrowheads mark lysosome-rich and deficient zones within a leader cell. **E**) Time stamps representing the lysosome-rich region (marked by green arrowhead in D) and lysosome-deficient region (marked by yellow arrowhead in D). Lamellipodial growth is observed in the lysosome-rich region, whereas lamellipodium retracts in the lysosome-deficient region. Scale bars, 10 µm for all sub-figures.

### Lysosome movement on microtubule tracks drives its peripheral localization and accumulation

Next, we asked whether the peripheral accumulation of lysosomes in the marginal cells, including the leader cells, was a result of increased lysosome biogenesis or of an escalated anterograde movement of the organelle. Considering that nuclear localization of Transcription Factor EB (TFEB) is an indicator of enhanced lysosome biogenesis^39, 40^, we examined TFEB localization. However, we did not observe TFEB accumulation in the nuclei of marginal cells at T-120 mins post confinement lift-off (Supplementary fig. 3A). We further quantified the number of lysosomes in the marginal and follower cells, before and during migration and did not observe any significant change (Supplementary fig. 3B). These results eliminated the possibility of *de novo* lysosome biogenesis and pointed towards the polarized transport of lysosomes in leader cells. Previous studies have shown that microtubules are the tracks for bi-directional molecular motor-mediated transport of lysosomes^41^. First, to gain insight into the organization of microtubules during collective epithelial migration, we performed a gap-closure assay with both MDCK and EPH4 monolayers. We allowed the cells were allowed to migrate for 0 and 120 minute and then fixed and immunostained them for microtubules with anti-Alpha-tubulin and lysosomes with anti-LAMP1. Microtubules show peripheral accumulation in marginal cells like that observed for lysosomes in both MDCK and EPH4 cells (Fig. 3A and Supplementary fig. 3C). Similar to computing the lysosomal POF, we also computed the POF (peripheral occupancy fraction) of microtubules to measure the microtubule accumulation at the cell periphery. We calculated the POF of microtubules by taking the ratio of the intensity of microtubules in the defined ROI to the total intensity of microtubules in the cell of interest. This analysis revealed a significant increase in POF of microtubules at T-2hrs post confinement lift-off like that was observed for lysosomes (Fig. 3B). Since lysosome movement is largely dependent on microtubule tracks, we envisaged that disrupting microtubules would abolish the dynamic peripheral accumulation of lysosomes. We, therefore, disrupted the microtubules by using nocodazole at a sub-optimal concentration (10 µM) that did not severely affect the collective migration of MDCK cells. Nocodazole treatment led to the loss of peripherally localized lysosomes (Fig. 3C, *middle panel*), and at the same time, we observed a reduced emergence of leader cells upon nocodazole treatment (Fig. 3D). Microtubules are dynamic in nature and constantly undergo cycles of assembly and disassembly at the plus end^42^. Taxol or Paclitaxel can stabilize the growing end of microtubules, thus altering their dynamics. We treated cells with Taxol again at a sub-optimal concentration (14 µM) that did not severely affect the collective migration of MDCK cells and observed a lysosome distribution pattern in this case. In this scenario, lysosome accumulation at the cell periphery abrogated, and they spread out evenly throughout the cytoplasm of marginal cells (Fig. 3C, lower panel). Taxol treatment also led to a significant drop in leader emergence (Fig. 3D). We quantified POF for lysosomes in control, nocodazole-, and taxol-treated cells and found a significant drop in the peripheral accumulation of lysosomes in the drug-treated cells as compared to the control (Fig. 3E). These results indicated that lysosome distribution depended on microtubule arrangement. Finally, we observed a temporally congruent polarization pattern for microtubules and lysosomes in MDCK cells, which were allowed to migrate for 0, 15, 30, 60, 120, and 240 mins post confinement lift-off (Supplementary fig. 3D).

**Figure 3:**
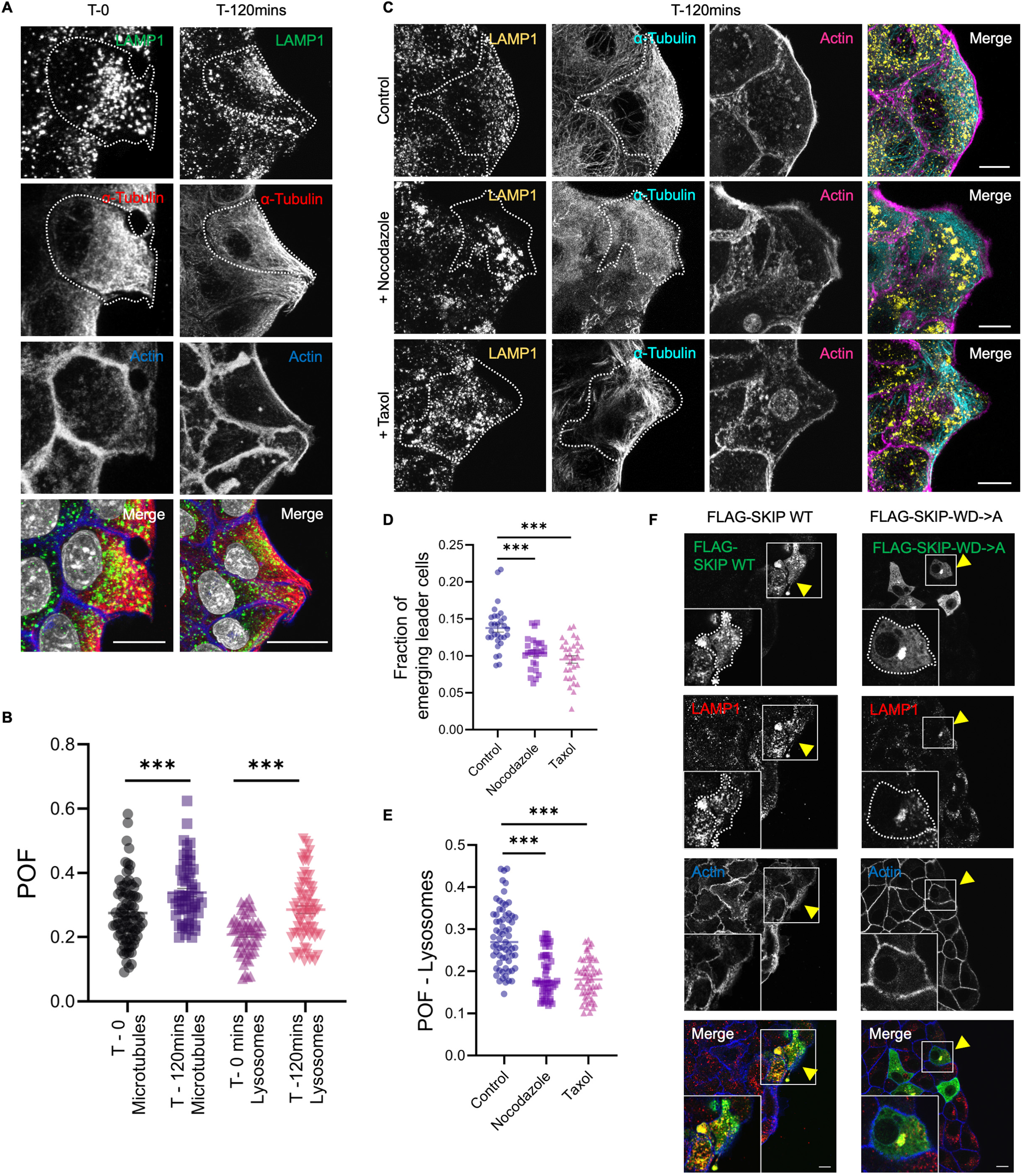
Lysosomes move on microtubules and accumulate to cell periphery in collectively migrating epithelial cells. **A**) Representative confocal micrographs of collectively migrating MDCK cells, fixed at T-0 and T-120mins post confinement lift-off and immunostained for lysosomes (Anti-LAMP1) and tubulin (Anti-α tubulin) and actin labelled with Alexa-fluorophore conjugated phalloidin. **B**) Scatter plot showing POF for microtubules and lysosomes, calculated using respective lysosome and tubulin staining intensity at T-0 and T-120mins for all cells at the margin (n=3, 20 cells per experiment). **C**) MDCK cells treated with sub-optimal concentrations of Nocodazole and Taxol were allowed to migrate for T- 120mins post confinement lift-off, PFA fixed and immunostained for lysosomes (Anti-LAMP1), tubulin (Anti-α tubulin) and actin labelled with Alexa-fluorophore conjugated phalloidin. **D**) Scatter plot showing the fraction of leader cell emergence in control, nocodazole and taxol-treated cells. Images were acquired for all treatments to cover the complete length of the margins of MDCK cells migrating collectively post-confinement lift-off (10 frames per treatment per experiment). The fraction of leader emergence was determined by taking the ratio of the number of cells showing leader cell features (lamellipodia and enlarged cell size) to the total cell number on the margin. The significance was calculated using one-way ANOVA, comparing Nocodazole and Taxol treatments with Control. **E**) Scatter plot showing POF of lysosomes in control, nocodazole and taxol-treated cells. Statistical analysis was done using one-way ANOVA comparing Nocodazole and Taxol treatments with Control (n=3; 20 cells per treatment per experiment were analysed). **F**) Confocal micrographs of MDCK cells transiently transfected with either FLAG-tagged SKIP WT or SKIP WD->A mutant, migrated for T-120mins post confinement lift-off, fixed and immunostained with anti-LAMP1 and Alexa-fluorophore conjugated phalloidin to label actin. The yellow arrowheads mark the transfected cells. Lysosomes are peripherally localized in **F**) *top panel* and juxtaposed to the nucleus in **F**) *bottom panel*. Scale bar, 10 µm for all sub-figures. ** and *** indicates p-value<0.01 and 0.001 respectively.

We next investigated if cellular organelles such as late endosomes, early endosomes, endoplasmic reticulum, and mitochondria which also use microtubule tracks for their intracellular transport, show peripheral accumulation. We again allowed MDCK cells to migrate for 0- and 120-mins post confinement lift-off, fixed and immunostained for late-endosomes (Anti-Rab7), early endosomes (Anti-EEA1), endoplasmic reticulum (mCherry-Sec61 stable cell line), and mitochondria (Anti-Tom20) (Supplementary figs. 4A-E). We observed that none of these organelles shows enhanced peripheral localization as compared to lysosomes. We compared the POF for all the organelles (Supplementary fig. 4F) and only lysosomes showed maximum peripheral enrichment at 120 mins post confinement lift-off. These results indicated that peripheral lysosomal accumulation is specific and not a passive effect due to microtubule re-orientation during collective cell migration.

Downstream effectors of small GTPase Arl8b control the lysosome movement on the microtubule track. Among these proteins, lysosome adaptor protein SKIP recruits the plus-end-directed kinesin motor to the lysosome^43^. We, therefore, hypothesized that transfecting the cells with a kinesin-binding defective mutant of SKIP (SKIP WD->2XA) would provide a more specific disruption of lysosome dynamics, without affecting the microtubule structure. Subsequently, SKIP mutant-expressing cells showed reduced peripheral accumulation of lysosomes (Fig. 3F). Taken together, these results indicated that peripheral accumulation of lysosomes in marginal cells is the result of a microtubule-mediated dynamic localization process of lysosomes.

### Peripheral accumulation of lysosomes is necessary for leader cell emergence

Next, we sought to examine whether the peripheral accumulation of lysosomes is necessary for the emergence of leader cells. We subsequently hypothesized that if that was the case, preventing lysosomes from moving to the cell periphery would reduce the propensity of leader cell emergence. Conversely, forcing lysosomes to the cell periphery would increase this propensity. To test these hypotheses, we first treated the MDCK epithelial monolayer with either Acetate Ringer’s (AR) or Alkaline Ringer’s (AlkR) solution for 30 minutes before the initiation of migration and let the cells migrate in the presence of these solutions for 4 hours. AR solution promotes peripheral accumulation of lysosomes while AlkR solution promotes perinuclear accumulation^44^ (Supplementary fig. 5A). Using the time-lapse images recorded every 15 minutes, we analysed the number of leader cells emerging from migrating monolayers. AR-treated Cells showing predominantly peripheral lysosomes also displayed a significantly higher fraction of leader cell emergence than AlkR-treated cells with perinuclear lysosomes (Supplementary figs. 5B-E). Together these results suggested that the propensity of leader cell emergence indeed depends on lysosome localization.

However, AR and AlkR solutions impart global perturbation to lysosome distribution, which may have secondary effects on migration. Hence, to specifically modulate lysosome distribution in cells, we switched to a method that allowed us tweaking of lysosome localization in individual cells. To alter the localization of lysosomes in a cell-specific manner, we resorted to the Reversible Association of Motor Protein (RAMP) system^45^. In this system, the interaction between streptavidin-bound to motor protein and streptavidin-binding protein (SBP) fused to an organelle-localizing protein drives the preferential localization of the target organelle. To this end, we first generated stable cell lines expressing streptavidin-bound to motor protein, mCh-Kif5b*-Strep or Strep-Kifc1*-mCh (Supplementary figs. 5F-G, mCh: mCherry). mCh-Kif5b*-Strep generates plus-end directed movement accumulating the target organelle at the cell periphery, while Strep-Kifc1*-mCh generates minus-end directed movement accumulating the target organelle near the cell nucleus (Fig. 4A and 4B, respectively). Further, we transiently transfected these cells with LAMP-1-SBP-GFP, targeting lysosomes, and seeded them in the migration chamber. By the interaction between Streptavidin and SBP, lysosomes were either localized to the cell periphery in mch-Kif5b*-Strep-expressing cells or the perinuclear region in Strep-Kifc1*-mCh-expressing cells (Figs. 4A-B). Using fluorescence time-lapse imaging, we then followed the trajectories of the LAMP1-SBP-GFP expressing cells at the migration front (Figs. 4C and 4D and Supplementary videos 3 and 4). Interestingly, the LAMP1-SBP-GFP-expressing cells in mCh-Kif5b*-Strep background, which had peripherally localized lysosomes (Fig. 4A and 4E), showed a significantly increased propensity of leader cell formation than the LAMP1-SBP-GFP-expressing cells in Strep-Kifc1*-mCh background (Fig. 4B and 4F), which had lysosomes mostly localized to the perinuclear region. Next, we calculated the probability of leader emergence in the two scenarios by dividing the number of co-transfected cells at the margin with lamellipodia (identified as leader cells) to the total number of co-transfected cells at the migrating edge. The probability of cells with peripheral lysosomes becoming leaders was 55% ± 3% (mean ± standard deviation), while same the probability of cells with perinuclear lysosomes was 7% ± 2% (Fig. 4G). Relevantly, the baseline probability was 20%±5% in mCh-Kif5b*-Strep and 16% ± 2% in Strep-Kifc1*-mCh motor-only controls (Fig. 4G). Moreover, we found that cells co-expressing mCh-Kif5B*-strep and LAMP1-SBP-GFP possessed prominent lamellipodial structures and mature focal adhesions oriented perpendicular to the migrating front (Fig. 4E). In contrast, cells co-expressing Strep-KifC1*-mCh and LAMP1-SBP-GFP possessed peripheral actin cables and less prominent focal adhesions oriented parallel to the migrating front (Fig. 4F). We also analysed the cell movement of LAMP1-SBP-GFP co-expressed with either mCh-Kif5b*-Strep or Strep-KifC1*-mCh and compared it to the motor-only control cells. These results suggested that mCh-Kif5B*-strep expression and subsequent peripheral localization of lysosomes promoted leader-like cell morphology, while mCh-Kifc1*-Strep expression and subsequent perinuclear localization of lysosomes promoted follower-like cell morphology^25, 26^. Finally, we obtained similar results using EPH4 cells, where cells transiently transfected with mCh-Kif5B*-strep and LAMP1-SBP-GFP or Strep-KifC1*-mCh and LAMP1-SBP-GFP showed lamellipodium or actin cables, respectively (Supplementary figs. 5H-I). Noticeably, abrogating lysosomal association with kinesin motor using the kinesin binding defective mutant of SKIP also showed a similar phenotype of reduced lamellipodial formation and hence leader emergence when compared to cells expressing wildtype SKIP (Fig. 3F). In summary, we were able to regulate the propensity of leader cell emergence at the migrating front by altering lysosome localization. Collectively, these results suggested that peripheral localization of lysosomes is both necessary and sufficient for leader cell emergence.

**Figure 4:**
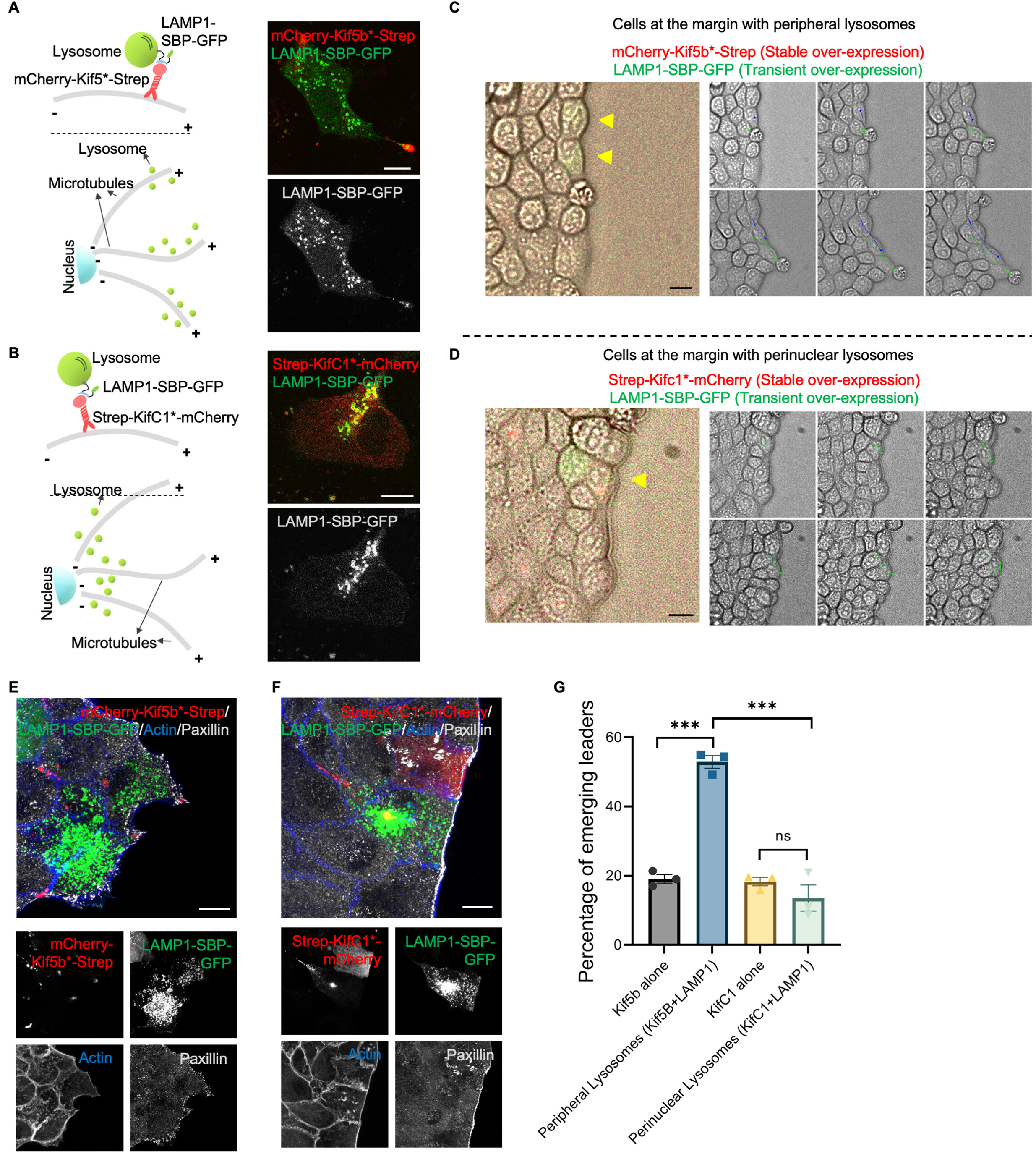
Peripheral lysosomal bias leads to enhanced leader cell emergence in collectively migrating epithelia. **A-B**) Schematic and representative confocal micrographs of the molecular basis and lysosome positioning outcomes using the RAMP (Reversible Association of Motor Protein) system. **C**) Representative fluorescence/bright-field time-lapse imaging snapshots of MDCK cells expressing mCherry-Kif5B*-Strep transiently transfected with LAMP1-SBP-GFP. Yellow arrowheads mark cells with co-expression of the two plasmids. **D**) Representative time-lapse imaging snapshots of MDCK cells expressing Strep-KifC1*-mCherry transiently transfected with LAMP1-SBP-GFP. Yellow arrowheads mark cells with co-expression of the two plasmids. **E-F**) MDCK cells co-expressing either mCherry-Kif5B*-Strep (**E**) or Strep-KifC1*-mCherry (**F**) and LAMP1-SBP-GFP were fixed 120 mins post-confinement lift-off and stained for focal adhesion protein using Anti-Paxillin and actin using Alexa-fluorophore conjugated phalloidin, further imaged using a confocal microscope. Cells with peripheral lysosomes (**E**) form extended lamellipodia, and cells with perinuclear lysosomes have actin cables (**F**). **G**) Graph representing the percentage of leader cells emerging in case of cells co-expressing mCherry-Kif5B*-Strep or Strep-KifC1*-mCherry and LAMP1- SBP-GFP, allowed to migrate for 240 mins, calculated as a fraction of co-transfected cells at the margin showing lamellipodial formation to the total co-transfected cells at the migration margin (n=3, ∼30 co-transfected cells per transfection combination per experiment were counted). The percentage of leader cell emergence was calculated in motor-only expressing cells as control. Scale bar, 10 µm for all sub-figures. *** signifies p-value < 0.001 and *n.s* is not significant.

### Forces generated by actomyosin contractility alter lysosome localization

In a cohesively migrating epithelium, leader cells emerge through an active mechanical process, where forces from cell-cell and cell-ECM interactions play critical roles^4, 6, 10, 46^. We, therefore, asked whether the peripheral accumulation of lysosomes in leader cells might be mechanosensitive. To test this possibility, we reduced the actomyosin contractility by using a non-muscle myosin II inhibitor, blebbistatin and a Rho-associated protein kinase inhibitor, Y-27632. We used sub-optimal concentrations of these drugs^25^ so that the cells were able to migrate (Fig. 5A). We pre-treated the cells for 30 minutes with the desired concentration of these inhibitors. We then lifted off the confinement and allowed the cells to migrate in a media containing these inhibitors. Subsequently, we observed a reduced accumulation of lysosomes at the cell periphery upon drug treatment (Fig. 5B, Supplementary video 5). In addition, there was no selective emergence of leader cells at the migrating front (Fig.5A). Time-lapse data of cells treated with actomyosin inhibitors revealed that the dynamics and speed of lysosomes were altered upon treatment (Supplementary figs. 6A-D). Lysosomes in cells treated with actomyosin inhibitors showed enhanced lysosomal tubular extensions (Supplementary figs. 6B-C). Together these results show that lysosome dynamics and localization is sensitive to the forces generated by actomyosin contractility. This set of experiments provided the first evidence supporting the mechano-sensitivity of lysosome localization during collective cell migration.

**Figure 5:**
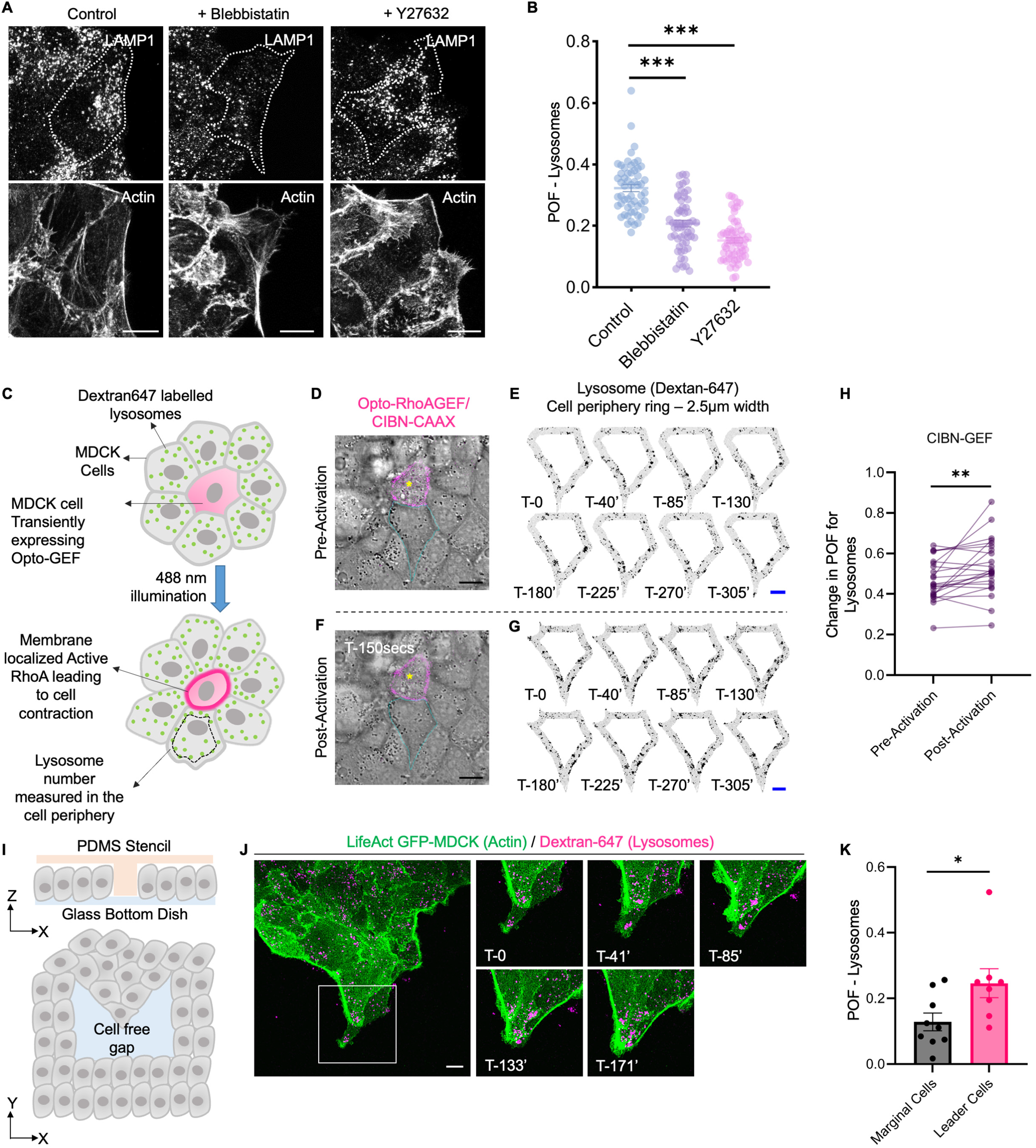
Actomyosin contractility drives peripheral localization of lysosomes. **A**) Control (DMSO), Blebbistatin (5µM), and Y27632 (30µM) treated MDCK cells were allowed to migrate for 120mins, fixed, and stained for lysosomes (anti-LAMP1) and actin (Alexa-fluorophore conjugated Phalloidin). Images shown were acquired on a confocal microscope. **B**) Graph representing POF of lysosomes for Control, Blebbistatin, and Y27632 treated MDCK cells (n=3; 25 cells per treatment per experiment). **C**) Diagrammatic representation of optogenetic control of Opto-RhoAGEF which allows manipulation of cell contractility and epithelial monolayer force landscape. Changes in lysosome distribution were recorded in cells juxta-positioned to the transfected and optogenetically altered cell. **D**) Live imaging snapshot of cells with Dextran-647-labelled lysosomes and expressing Opto-RhoAGEF. The juxta-positioned cell is marked by a cyan boundary. **E**) Pre-activation grayscale inverted image montage of 2.5µm wide cell periphery showing lysosome punctae in the cell marked by cyan outline in D). **F-G**) Post-activation with 488nm laser illumination, lysosome distribution was observed in the neighbouring cell marked by cyan outline. Lysosome punctae can be observed in 2.5µm cell periphery in the representative grayscale inverted image montage. **H**) Fraction of peripheral lysosome was calculated by dividing lysosome intensity in 2.5µm cell periphery ring to the total lysosomal intensity in pre-activated and post activated condition in cell juxta positioned to optogenetically altered cells. Dot plot showing peripheral lysosome fraction in cells juxta-positioned to Opto-RhoAGEF/CIBN-CAAX expressing cells under pre-activated and post-activated conditions. (n=3; 25 cells were analysed). **I**) Schematic representing micropatterned single beak mold used for defining leader cell-like characteristics in select cells. **J**) Dynamics of dextran-647-labelled lysosomes were observed in LifeAct-GFP MDCK cells seeded in micropatterned chambers. Cells were allowed to migrate for up to 180 mins post lifting off the confinement. **K**) Graph representing POF of lysosomes calculated for non-leader marginal and leader cells (n=3; 15 cells analysed per condition). Scale bar, 10 µm for all sub-figures; *, **, *** and **** signifies p-value<0.05, 0.01, <0.001 and <0.0001 respectively.

To test the connection between the actomyosin contractility and the peripheral accumulation of lysosomes further, we next performed two independent sets of experiments. Previous studies had shown that increased traction forces and tensile stresses among the follower cells preceded leader cell emergence^25^. This observation is connected to the fundamental premise of contact inhibition of locomotion (CIL), where one can imagine a leader-follower doublet to be a connected pair of cells experiencing a force perpendicular to the cell-cell contact^47, 48^ (Supplementary fig. 7A). Therefore, increased tension in one cell will trigger the formation of lamellipodium in the other cell, away from the first cell (Supplementary fig. 7A). Hence, we hypothesized that in such a cell-pair system, lysosomes would localize towards the cell periphery, away from the cell-cell interface. Moreover, this localization would depend on the actomyosin contractility. To test this hypothesis, we plated MDCK cells at a low density to allow them to form cell pairs and subjected them to overnight treatment with blebbistatin to reduce the actomyosin contractility (Supplementary fig. 7B). As compared to the control, polarized localization of lysosome, away from the cell-cell interface, was lost in blebbistatin-treated cells, and lysosomes dispersed throughout the cell body (Supplementary fig 7B). To test whether lysosome polarization could be restored upon the revival of actomyosin contractility, we performed subsequent drug washout experiments. Cells maintained under drug-free conditions for 4 hours partially regained the lysosome polarization (Supplementary fig. 7B). These results indicated that the force experienced at the cell-cell interface of a leader-follower pair is crucial for lysosome localization.

We further examined whether locally enhanced cellular forces within an epithelial monolayer might be sufficient to induce lysosome polarization away from the interface. To this end, we used an Opto-GEF-based strategy^49^ to enhance the actomyosin contractility of a specific cell within the monolayer (Fig. 5C). This experiment involved blue light-mediated photo-stimulation of cryptochrome 2 (CRY2)-bound catalytic domain of a RhoA-activator, ARHGEF11. CIBN, which is one of the binding partners of CRY2, was engineered to localize either at the plasma membrane or mitochondria. Upon photo-stimulation, CRY2-CIBN association triggered plasma membrane (Fig. 5C and Supplementary video 6) or mitochondrial localization (Supplementary Fig. 7C and Supplementary video 7) of the RhoA-activator. When localized to the plasma membrane, RhoA-activator increased the actomyosin contractility^49^. We used the mitochondrial localization of the RhoA-activator as the negative control for this experiment. Subsequently, we hypothesized that enhanced contractility of a photo-stimulated cell would polarize the lysosomes in the neighbouring cells, away from the former (Fig. 5C). As expected, upon photo-stimulation, lysosomes in the cells adjacent to the excited cell moved towards the cell periphery, away from the stimulated cell (Figs. 5D-H). But, in the negative control, where RhoA-activator localized to mitochondria, lysosomes in the cells adjacent to the stimulated cell did not undergo any such changes in localization (Supplementary figs. 7D-E). These two sets of experiments revealed how under the framework of CIL, mechanical forces transmitted through the cell-cell interface polarized lysosomes away from the interface. Lysosomes, therefore, appear as the missing cellular connection between increased forces among the follower cells and the emergence of leader cells.

Finally, we hypothesized that if the mechanical forces regulated peripheral lysosome localization during the emergence of leader cells, then any cue that spatially biases leader cell emergence should also spatially bias the peripheral localization of lysosomes. One such cue is wound geometry. Having highly positive-curved beaks at the migration front leads to the emergence of leader cells at the tip of the beak, due to strong force concentration at those points^50–52^. Would such beaks show enhanced peripheral localization of lysosomes? To answer this question, we used a micropatterning approach to fabricate silicone stencils made of polydimethylsiloxane (PDMS). These stencils would create wounds with a single beak to allow one or two cells to have the geometrical cue to become the leader cells (Fig. 5I and Supplementary fig. 7F). We then seeded cells in these micropatterned chambers, labelled lysosomes using Dextran-647, lifted off the confinement, and performed live-cell imaging. The cells seeded in these micropatterned chambers moved collectively and covered the gap (Supplementary fig. 7G and Supplementary video 8). Upon lifting the stencil, we indeed observed enhanced lysosome accumulation at the tip of these beaks as compared to their sides (Figs. 5J-K and Supplementary video 9). At the same time, lamellipodial structures emerged out of the beak region while actin fibres ran along both sides (Fig. 5J). These results suggested that the spatial concentration of mechanical force in the epithelial monolayer correlates with the enhanced peripheral localization of lysosomes, supporting the premise of mechanosensitive localization of lysosomes during leader cell emergence. It also strengthened the correlation (Fig. 2D) between peripheral localization of lysosomes and lamellipodia formation through the modification of the actin cytoskeleton.

### Inactive Rac1 GTPase localizes to lysosomal membranes

We then asked how the peripheral localization of lysosomes promotes the lamellipodia formation by locally modifying the actin cytoskeleton. In migrating epithelial cells, actin dynamics depend on the localization and activation of small Rho GTPases, including Rho, Rac1, and Cdc42^3, 11, 26, 53^. These proteins switch between GTP-bound active state and GDP-bound inactive state. To identify if any form of these three small Rho GTPases interacted with lysosomes, we conducted a screen using the constitutively active (CA) and dominant-negative (DN) forms of these GTPases (Figs. 6A-C and Supplementary figs. 8A-D). These proteins were also tagged with a fluorescent protein for visualization. Unfortunately, antibodies claiming to detect the endogenous GTP or GDP were not reliable, and a pull-down approach will not give us the cell-specific information in a dynamic environment such as during the collective cell migration. Hence, we used the CA and DN forms of the small RhoGTPases for our screening, which represented the active (mimic of GTP-bound molecules) and inactive (mimic of GDP-bound molecules) states of the small Rho GTPases, respectively. We transfected MDCK cells with one of the forms, grew these cells under confinement, and lifted off the confinement to allow them to migrate for 4 hours. We then fixed and immunostained these cells to visualize lysosomes (Figs. 6A-C and Supplementary figs. 8A-D). To our interest, we observed that out of these six proteins, only the dominant-negative form of Rac1 GTPase (Rac1-DN) colocalized with the lysosomal marker LAMP1 (Fig. 6C). Other Rho family GTPases did not show any appreciable colocalization with lysosomes (Supplementary figs. 8A-D). Also, the wildtype (Rac1-WT) and Rac1-CA were primarily plasma membrane-localized whereas Rac1-DN was distributed on lysosomes and at the plasma membrane in both MDCK and EPH4 epithelial cells (Figs. 6A-C and Supplementary figs. 8E-G). Mander’s coefficient analysis between Rac1-WT, Rac1-CA, and Rac1-DN with the lysosomal membrane marker LAMP1 confirmed this localization pattern (Fig. 6D and Supplementary fig. 8H). We corroborated our finding by using super resolution by optical pixel reassignment (SoRa) microscopy to study the co-localization of Rac1DN with dextran-647 labelled lysosomes (Fig. 6E). We live-imaged MDCK cells transiently transfected with GFP-Rac1-DN and labelled with Dextran-647 and observed GFP-Rac1-DN encircling Dextran-647 labelled lysosomes (Fig. 6E). With these results, we wondered whether lysosomes are capable of scavenging Rac1-DN. To reveal the dynamic association between lysosomes and Rac1-CA or Rac1-DN, we used a photoconversion strategy. In this strategy, we expressed either mEos-Rac1-DN or mEos-Rac1-CA in MDCK cells having lysosomes labelled with Dextran-647. Upon photoconversion using 405 nm illumination, the emission spectrum of mEos fluorescent protein shifted from green to red, allowing us to visualize the dynamics of the protein of interest. In this experiment, we allowed mEos-Rac1-DN or mEos-Rac1-CA transfected cells to migrate, and during migration, we photoconverted the colour of mEos proteins near the growing lamellipodial protrusions using a 405 nm source. We then tracked the photoconverted proteins for their redistribution within the cell (Figs. 6F-J). As depicted by kymographs, live-tracking revealed puncta formation events for photoconverted mEos-Rac1-DN. Interestingly, these punctate structures merged with lysosomes positioned nearby, showing significant colocalization between the two (Fig. 6G and Supplementary video 10). In contrast, photoconverted mEos-Rac1-CA diffused throughout the plasma membrane, and puncta formation was negligible in this case (Fig. 6I and Supplementary video 11). Collectively, these results indicated lysosomes had a specific affinity for scavenging only the inactive form of Rac1, which piggybacked on the lysosomal membrane and moved away from the plasma membrane (Figs. 6E-J). These results also suggested that the presence of lysosomes at the cell periphery should lead to an increased Rac1 activity, leading to branched actin polymerization and lamellipodium formation. To test this prediction, we measured the peripheral Rac1-activity with a FRET-based Rac1 sensor^53^ in the presence or absence of AR solution, which promotes peripheral accumulation of lysosomes^44^. This experiment yielded two key results. First, the basal level Rac1 activity in quiescent cells (T= 0) increased upon AR treatment (Figs. 6K-M). Second, although there was an increase in Rac1 activity during the cell migration even in the absence of AR-treatment, upon AR-treatment, we observed a further increase in Rac1 activity (Figs. 6L). In addition, upon AR treatment, many migrating cells showed unusual projections throughout the cell periphery (Fig. 6L, *right panel)*. Taken together, these results suggested that the peripherally localized lysosomes promoted Rac1 activity, thus linking the force-induced peripheral accumulation of lysosomes to lamellipodium formation in the emergence of leader cells during collective cell migration (Fig. 7)

**Figure 6:**
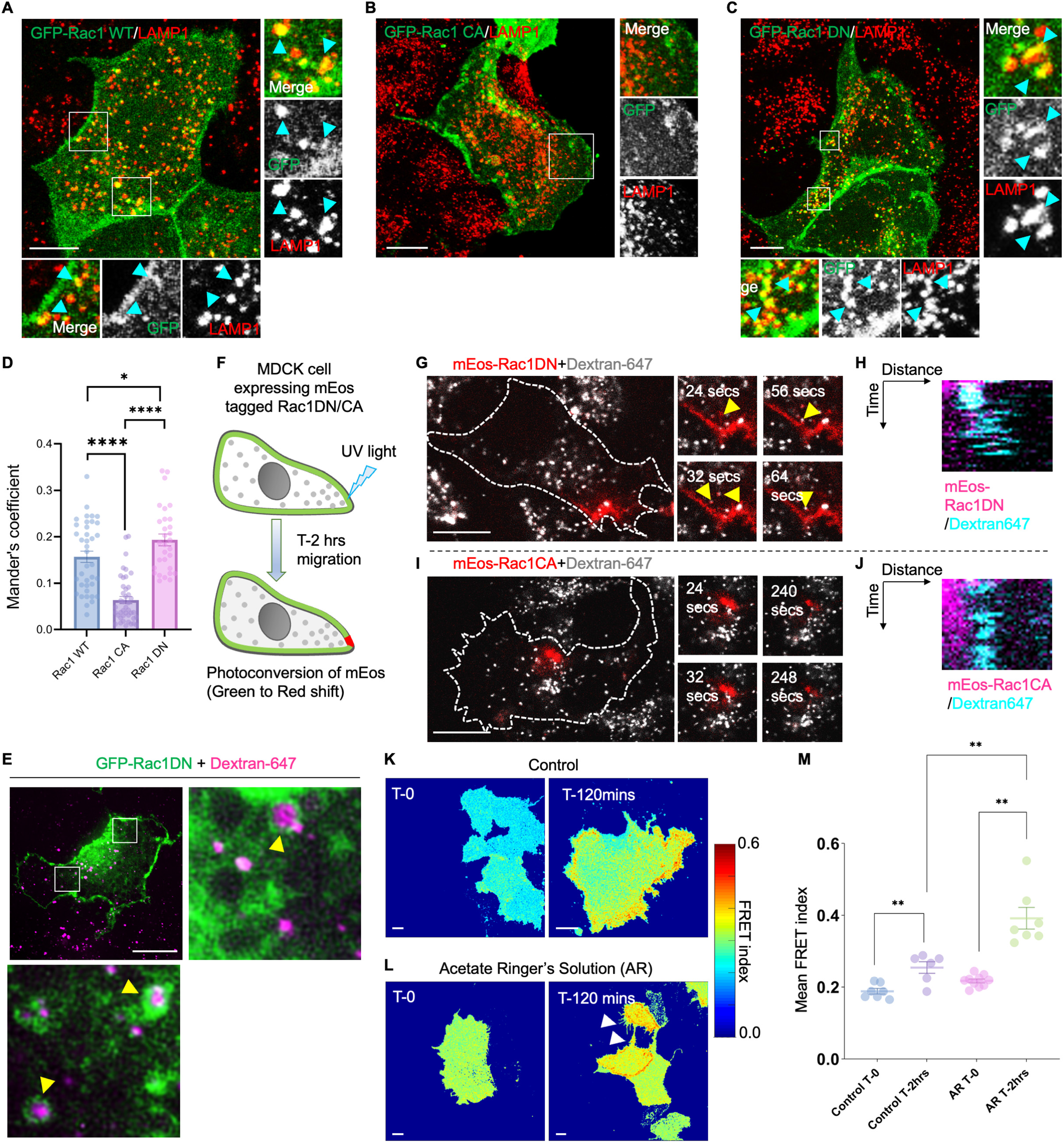
Peripheral lysosomes sequester inactive Rac1 from sites of growing lamellipodia. **A-C**) Representative confocal micrographs of MDCK cells transiently transfected with GFP-Rac1 WT, DN (dominant-negative; inactive form), and CA (constitutive active: active form), respectively, and allowed to migrate for 120 mins, followed by fixation and immunostaining for lysosomal markers LAMP1. **D**) Colocalization of Rac1 WT, DN, and CA forms were assessed with LAMP1 by measuring Mander’s coefficient (n=3, 15 cells per transfection per experiment were analysed; data represented as mean±sem). **E**) SoRa super-resolution images showing GFP-Rac1DN recruitment onto Dextran-647-labelled lysosomes in MDCK cells. **F**) Schematic depicting the mEOS-Rac1DN/CA transiently transfected in MDCK cells and photoconverted using UV light which leads to a green to red shift in emission, thus allowing tracking of their localization dynamics. **G**) Photoconversion of mEos-Rac1DN to track its localization dynamics. MDCK cells transiently transfected with mEos-Rac1DN and dextran-647 labelled lysosomes were allowed to migrate for 60 mins before photoconversion to initiate lamellipodia formation. Dynamics of Rac1DN puncta formation and trajectory were followed for photoconverted mEos-Rac1DN. Yellow arrowheads mark emerging Rac1DN puncta and their localization to dextran-647 labelled lysosomes. **H**) Kymograph representing the colocalization dynamics of lysosomes and photoconverted mEos-Rac1DN. **I**) Photoconversion of mEos-Rac1CA to track its localization dynamics. Similar to mEos-Rac1DN, mEos-Rac1CA was photoconverted post 60mins of migration, and its dynamics and colocalization with dextran-647-labelled lysosomes were recorded. **J**) Kymograph representing distribution dynamics of lysosomes and photoconverted mEos-Rac1CA. No overlap was observed for lysosomes and mEos-Rac1CA. **K-M**) Representative heat maps of FRET-based Rac1 activity sensor for Control, Acetate Ringer’s and Alkaline Ringer’s treated MDCK cells. White arrowheads mark multiple cell protrusions observed upon Acetate Ringer’s treatment. **N**) Mean FRET index was calculated for Control, Acetate Ringer’s and Alkaline Ringer’s treated cells at T-0 and T-120mins and represented as a scatter column graph. Scale bar, 10 µm for all sub-figures; *, ** and **** signifies p-value<0.05, 0.01 and 0.0001 respectively.

**Figure 7:**
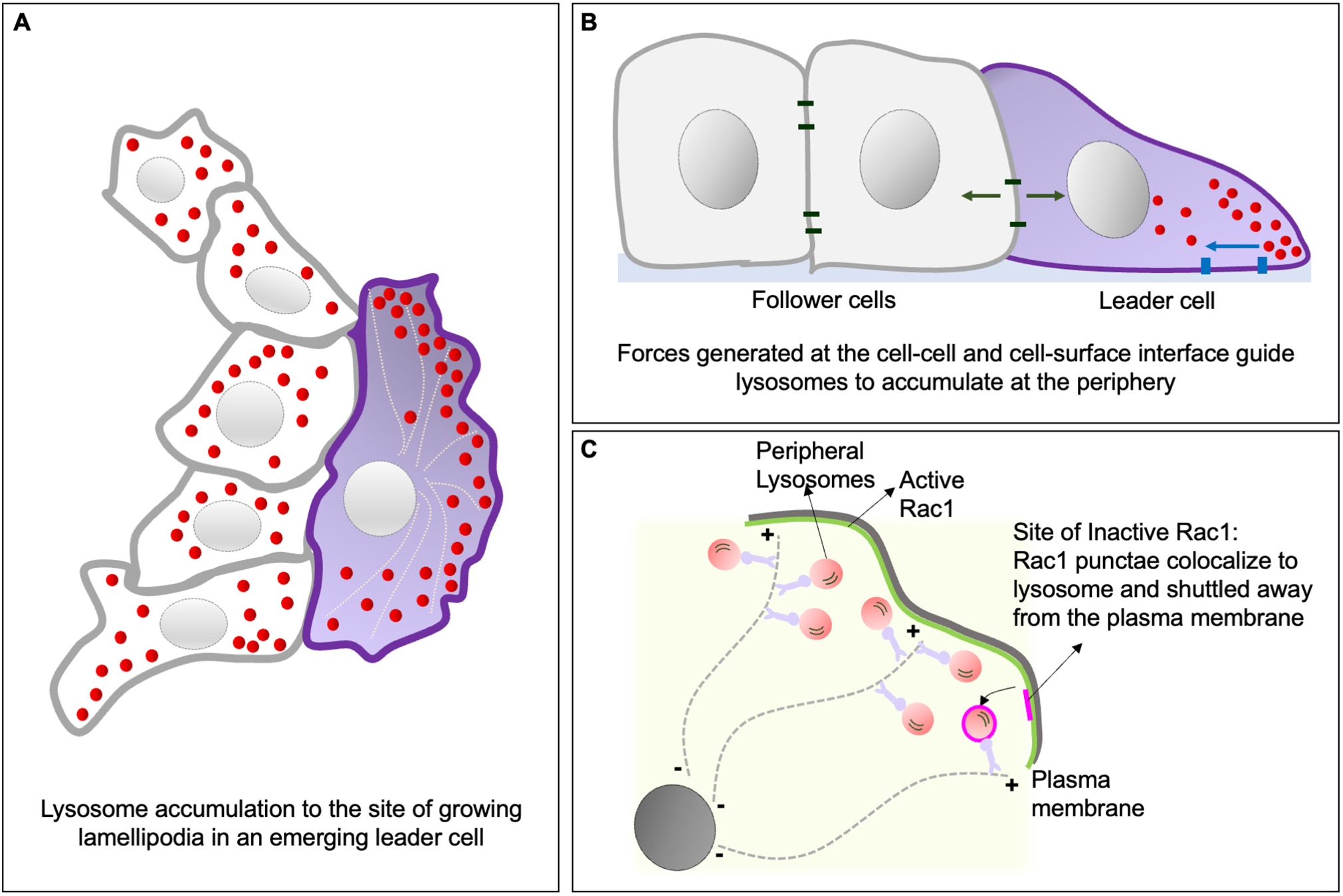
Role of lysosomes in regulating leader cell emergence. As epithelial cells start migrating collectively, lysosomes in the cells at the wound margin start accumulating at the cell periphery in the direction of migration. Two hours into migration, leader cells start emerging from the cell collective. As some cells emerge as leaders, lysosomes in these cells show a significantly higher accumulation of lysosomes at the periphery. This peripheral pool of lysosomes facilitates lamellipodial extension in leader cells (**A**). Disruption of the peripheral lysosomal pool severely reduces leader cell emergence. The cue for peripheral distribution of lysosomes comes from the mechanical forces experienced by an emerging leader cell. Forces generated at the cell-cell and cell-surface interface are translated to changes in actomyosin contractility within a leader, which guides the peripheral accumulation of lysosomes (**B**). Lysosomes in leader cells regulate the growth of lamellipodium by shuttling the inactive Rac1 away from the plasma membrane, thus shifting the balance towards active Rac1 which further regulates the formation of actin crosslinks required for lamellipodial extension (**C**).

## DISCUSSION

Ever since Edward Ruth published his pioneering study on epithelial wound closure using the fragments of frog skin^54^, the ubiquity and importance of leader cells in collective cell migration have become apparent in various physiological contexts^1–3, 12, 17^. Yet, for more than a century, it has remained intriguing how these special cells, with their characteristic lamellipodial protrusions, emerge from a seemingly homogeneous population. Once the free edge is generated by wounding, cells at the wound margin experience an asymmetry. While cells within the epithelium are encircled by neighbouring cells from all sides, cells at the migration front have at least one cell-free edge. Consequently, the geometric, physical, and molecular asymmetry arising at this free edge creates a spontaneous cue for polarized activation of small Rho GTPases, whose activity would otherwise be restricted by cell-cell junctions. In fact, asymmetric spatiotemporal regulation of Rho GTPases, including Rac1, is central to forming actin projections and stress fibres during collective cell migration, especially in the emergence of leader cells^3^. However, this asymmetry is equally likely to generate lamellipodial structures in all cells at the migration front, and it is not obvious at what level the selection of a leader cell is decided. Previous experiments have shown that leader cell emergence is not an entirely stochastic biochemical event, as conventionally believed. Instead, the localization of highly tensile peaks in the stress landscape determines the precise location of the leaders. These experiments have also suggested that enhanced tensions at the cell-cell junction promote the formation of lamellipodial protrusions away from the leader-follower interface, towards the cell-free space, in accordance with the fundamental principles of contact inhibition of locomotion^47, 48^ (Supplementary fig. 7A). However, the connection between this biophysical driving force and the eventual activation of Rac1 leading to lamellipodia formation remained elusive. It was not clear how these biophysical and biochemical paradigms could be unified under a single mechanistic framework.

The present study identifies lysosomes as an intracellular platform linking mechanical and biochemical signals towards the regulation of leader cell emergence. At one end, lysosomes respond to changes in the force landscape of the emerging leaders by localizing to the cell periphery (Fig. 7). At the other, they transiently colocalize with the inactive form of small GTPase Rac1. This specific lysosome-Rac1 interaction supports lamellipodia formation by shuttling the inactive form of Rac1 away from the plasma membrane. This process shifts the balance in favour of actin crosslinking to form migratory lamellipodial structures (Fig. 5). Thus, lysosomes sense the local force field and regulate lamellipodia formation by modulating the Rac1 signalling. This forms a positive feedback loop where cell-cell forces determine a leader cell, in which the lysosome polarization enhancement leads to the regulation of Rac1 signalling in these emerging leaders. Lysosomes are dynamic organelles that move bidirectionally on microtubules. Previous studies have shown that the position of lysosomes is critical for their function^33^. Here, we elucidate how lysosome relocalization inside a cell at the migrating front determines the probability of that cell emerging as a leader cell, by a novel mechanism involving lysosome-Rac1 interaction. Re-localization of lysosomes to the cell periphery is specific as other organelles such as late endosomes, early endosomes, endoplasmic reticulum, and mitochondria do not show significant accumulation during collective cell migration. Next, it will be interesting to probe how lysosomes influence the dynamics of other factors, including p53^2^ and Dll4^28^, in the context of leader cell emergence. Also, the underlying mechanism that makes lysosome mechanosensitive, needs to be elucidated. It is highly possible mechanosensitive peripheral accumulation of lysosomes is an outcome of the mechanosensitive reorientation of microtubules during collective cell migration.

Relevantly, Rac1 molecules organize into nanoclusters at the plasma membrane and form a gradient of spatial signalling. Rac1 is known to stay active for a few minutes in migrating cells^55, 56^. We speculate that lysosomes shuttling the inactive Rac1 away from the site of growing lamellipodia are essential for maintaining greater pools of active Rac1, which can further recruit actin remodelling proteins such as Arp2/3 and WAVE. A previous work^57^ has shown that early endosomes act as sites for Rac1 activation and transport activated Rac1 to the plasma membrane during single-cell migration. It will be interesting to explore the crosstalk between lysosomes and early endosomes to further characterize the signalling pathways underlying leader cell formation during epithelial migration^11^. Finally, the coherent observation of lysosome accumulation in the leader cells of mammalian epithelial monolayer and *Drosophila* epidermis indicates that the process we discovered could be evolutionarily conserved in epithelial wound healing. However, it remains unknown whether lysosome-Rac1 interaction is also critical for other kinds of single and collective cell migrations^4^, including neural crest cell migration in vertebrates^4^, border cell migration in *Drosophila*^11^, and collective migration of lateral line primordial cells in Zebrafish^58^. In addition, given that lysosomes play an important role in the continuous cross-layer migration and differentiation of keratinocytes within a mammalian epidermis^59^, the lysosome-Rac1 signalling axis might be critical for the development and maintenance of the multi-layered epidermal epithelium. Collectively, the results of our experiments and their implications place the functioning of lysosomes at a crucial junction of mechanochemical signals during collective cell migration and uniquely expand the scope of this organelle beyond its conventional role in the catabolism of cellular materials.

## MATERIALS AND METHODS

### Cell culture

Madin-Darby canine kidney (MDCK) and EPH4-Ev epithelial cells were used in this study. Tetracycline-resistant wild-type MDCK (MDCK-WT) cell line was a gift from Yasuyuki Fujita. MDCK cells were cultured in Dulbecco’s modified Eagle’s medium (DMEM) supplemented with GlutaMax (Gibco) with 5% foetal bovine serum (tetracycline-free FBS, Takara Bio) and 10 U mL^−1^ penicillin and 10 μg mL^−1^ streptomycin (Pen-Strep, Invitrogen) in an incubator maintained at 37°C and 5% CO_2_, unless mentioned otherwise. EPH4-Ev cells (ATCC; CRL-3063) were cultured in Dulbecco’s modified Eagle’s medium supplemented with GlutaMax (Gibco) with 10% foetal bovine serum (tetracycline-free FBS, Takara Bio) and 1.2 μg ml−1 puromycin (Gibco) in an incubator maintained at 37 °C and 5% CO2.

MDCK cells stably expressing mCh-Kif5b*-Strep and Strep-KifC1*-mCh (mCh: mCherry), cells were transfected with the respective plasmids using Lipofectamine 2000 (Invitrogen) following the instructions provided by the manufacturer. 16-18 hours post-transfection, cells were seeded via serial dilution in a 96-well plate, such that each well had a single cell. Forty-eight hours post-cell seeding, the cells were subjected to selection media (DMEM-GlutaMax plus 5% FBS) containing 400 μg mL^−1^ of Geneticin (Invitrogen). The growth of single cell-derived colonies was monitored over two weeks, following fluorescence confirmation. Colonies with a homogenous expression of the fluorescently-tagged protein of interest were expanded and further maintained in 100 μg mL^−1^ Genetic containing media. All transient transfections were done using Lipofectamine 2000 following following the instructions provided by the manufacturer.

Cell migration assays were performed using adhesive biocompatible silicone culture inserts from iBidi. 5 x 10^4^ cells suspended in an 80 μL cell culture medium were seeded in a culture insert stuck to glass-bottom dishes and incubated at 37°C in a 5% CO_2_ humidified chamber. After 18 hours of cell seeding, the cells formed a monolayer, and the culture insert was lifted to allow the cells to migrate. All migration experiments were carried out at 37°C and in a 5% CO_2_ humidified chamber (standalone or stage-top incubator).

### *Drosophila melanogaster* embryo microinjection wounding assay

For fixation and immunostaining, a previously described protocol was followed^60^. Briefly, Stage 15 *Drosophila melanogaster* embryos were collected on an apple juice plate. These embryos were then aligned and stuck to a double-sided tape on a glass slide. They were left at room temperature for 5 mins, followed by halocarbon oil application and subsequent wounding using glass needles mounted on an Eppendorf Femtojet system. At 15 mins post-wounding, halocarbon oil was drained from the slide and flushed with heptane to release the embryos from the double-sided tape. Next, 0.8 mL 4% Formaldehyde (in PBS) was added to a fresh tube and embryos were transferred with heptane to this tube, which was then placed on rotation in an orbital shaker for 25 mins. The aqueous solution was then removed from the bottom of the tube, and subsequently, heptane was removed too. To devitellinize the embryos, heptane was rapidly added followed by adding double to triple the volume of 90% ethanol and mixing it vigorously by vortexing. We then proceeded with embryos that settled down in the tube. The embryos were given 5 quick washes with 1X PBS and then blocked in PBS+0.3%Triton-X100+0.5%BSA (PBTB) for 30 minutes at room temperature. Primary antibody against LAMP (Abcam) was prepared in PBTB. Embryos were incubated overnight at 4°C followed by three washes done in PBS+0.3%TritonX-100 (PBT). Secondary antibody was prepared in PBTB and incubated for 2 hours at room temperature followed by 5 washes with PBT. DAPI and Phalloidin staining was done along with the secondary antibody. The embryos were mounted in Fluoromount (Sigma) on a glass slide and imaged using confocal microscopy.

### Mouse embryonic skin wounding

E14.5 - E16.5 mouse embryos were dissected out. The dorsal skin from these embryos was then excised. The skin was placed on a filter paper (Whatman filter, 8 μm pore size) with the epidermal side up. Using a scalpel/fine scissor, incision wounds were made on the epidermis. The filter paper along with the tissue was kept floating in a 6-well plate containing 2 mL of DMEM with 5% fetal bovine serum (FBS) and 1% PenStrep. After the desired incubation period, the tissue was fixed in 4% paraformaldehyde (PFA) overnight at 4°C. The tissue was washed in PBS and transferred to 90% ethanol in PBST (0.2% Triton X-100 in 1X PBS) overnight at -20°C. The tissue was then rehydrated using graded ethanol (75%, 50% and 25% in PBST). After two more washes in PBST, the tissue was blocked using 10% normal goat serum (NGS) in PBST for 2 hours. Primary antibodies were added and incubated overnight at 4°C. This was followed by subsequent PBST washes, after which secondary antibodies were added and incubated for 2 hours at room temperature. After washing, the tissue was mounted on a slide using Fluoroshield (Sigma Aldrich) mounting media. The slide was cured overnight and imaged using an upright confocal (Leica Stellaris).

### Immunofluorescence

Cells were fixed in 4% formaldehyde diluted in 1X PBS for 15 minutes at room temperature. For most antibody staining, cells were permeabilized with 0.25% TritionX-100 in PBS for 10 minutes at room temperature followed by three quick washes with 1X PBS to remove the detergent. Cells were treated with blocking/staining solution (0.1% TritonX-100in 1X PBS + 2% BSA) for 1 hour at room temperature, washed thrice with 1X PBS, and further incubated with primary antibody prepared in blocking solution for 2 hours at room temperature. Post-primary antibody incubation cells were washed thrice with 1X PBS and incubated with secondary antibody, DAPI, and Phalloidin dilutions prepared in the blocking/staining solution for 1 hour at room temperature. Finally, the samples were given 2 quick and 5 minutes wash with 1X PBS before proceeding for confocal microscopy.

For staining LAMP1 (Abcam), cells were incubated with a blocking solution (0.2% Saponin+5% BSA in 1X PBS) for 30 minutes at room temperature followed by three washes with 1X PBS. Cells were incubated with the primary antibody in staining solution (PBS+0.2%Saponin) for 2 hours at room temperature, washed thrice with 1X PBS and further incubated for 1 hour with secondary antibodies, DAPI (4′,6-diamidino-2-phenylindole, 1μg mL^−1^ in PBS) and Phalloidin made in staining solution. Cells were washed thrice with 1X PBS and imaged using confocal microscopy.

### Lysosome labelling for live cell imaging

Lysosomes of MDCK cells for live-cell imaging experiments were labelled with Alexa-fluor conjugated 10,000 MW Dextran (Dextran-647 from Invitrogen). Briefly, cells were incubated with Dextran-647 at a final concentration of 0.25mg ml^−1^ in complete media for 6 hours at 37 °C in a 5% CO2 humidified chamber. Media for cells was then replaced with complete media and incubated for another 12 hours to completely chase Dextran-647 to lysosomes.

### Confocal Microscopy

Fluorescence images were acquired using 60X oil objective (PlanApo N 60x Oil, N.A=1.42, Olympus) and 100X oil objective (UPlanSApo, 100X/1.40 oil) mounted on an Olympus IX83 inverted microscope equipped with a scanning laser confocal head (Olympus FV3000). Time-lapse images of live samples were done in the live-cell chamber provided with the microscopy setup.

Photoconversion studies were done using MDCK cells that had been transiently transfected with mEos-Rac1 DN or mEos-Rac1 CA. Transiently transfected cells were seeded in culture inserts and proceeded for lysosome labelling using Dextran-647 as described above. Cells were allowed to migrate for 60 minutes to allow for lamellipodia formation in migrating cells Stimulation was done on a point region of interest juxtaposed to the site of growing lamellipodia using a 405 nm laser at 2% intensity, looped over 25 times with a scan speed of 1000 μs/pixel. This was followed by LSM imaging of the green and red channels and Rac1 dynamics were tracked for 5 minutes.

The photoactivation experiment was carried out using a 60X oil objective (PlanApo N 60x Oil, NA=1.42, Olympus) mounted on an Olympus IX83 inverted microscope equipped with a scanning laser confocal head (Olympus FV3000), supported with a live-cell imaging setup. 20 mM HEPES (Gibco) was used to maintain CO2 levels. MDCK cells expressing *optoGEF-RhoA* and *CIBN-GFP-CAAX* or *optoGEF-RhoA* and *Mito-CIBN-GFP* (control)^49^, stained with lysosomal marker Dextran-647 were used for the experiment. Pulse activation (20 iterations) was performed in the selected rectangular ROIs (whole cell or cell-cell junction) using a 488 nm laser with 2.0 % laser intensity at a 4.0 μsec/pixel scan rate. This step was immediately followed by time-lapse (5 minutes, 10-sec intervals) LSM imaging in 561 nm and 647 nm channels, to visualize RhoA activation and monitor lysosomal movement, respectively. For super-resolution microscopy, images were obtained using a 100X oil objective (Nikon) on an inverted microscope equipped with a Yokogawa CSU-W1 (SoRa Disk) scanner.

### Förster resonance energy transfer (FRET)-based Rac1-activity measurements

FRET experiments for the Rac1 activity sensor were carried out in the live-cell confocal setup (Olympus FV3000). MDCK cells were first plated in a 24-well plate and transiently transfected with Rac1 sensor plasmids^53^. After 6 hours, cells were trypsinized and seeded in migration chambers and left overnight to settle and form confluent monolayers in the chamber. Cells were simultaneously labelled for lysosomes using Dextran-647. Control and AR treatment was given to cells as described in the Global perturbation of lysosome positioning using Ringer’s solution method section. The images were acquired for control and AR-treated cells just after lifting off the confinement (T-0) and post 2 hours of migration (T-2 hours). The exposure times for donor, acceptor, and FRET channels were kept constant. Images of Dextran-647 labelled lysosomes were also acquired. Each field yielded three 800 x 800-pixel images representing donor, acceptor, and FRET channels. Images were analysed using custom software written in MATLAB (Mathworks).

### Inhibition studies

For all inhibition studies, cells were pre-treated with the desired concentration of the inhibitor in opti-MEM for 30mins at 37°C in a 5% CO2 humidified incubator, before lifting the confinement/culture insert. During migration, cells were kept in complete media containing the inhibitor at a given concentration for the required time. Actomyosin contractility was altered using Blebbistatin (Sigma), a myosin inhibitor and Y27632 (Sigma), ROCK inhibitor at concentrations 5 μM and 30 μM respectively for MDCK cells. For microtubule disruption in MDCK cells, Nocodazole (Sigma) was used at 10 μM, and Taxol/Paclitaxel (Sigma) was used at 14 μM. Cells were fixed and immunostained for the desired antibodies post-inhibitor treatment.

### Global perturbation of lysosome positioning using Ringer’s solution

Lysosome positioning in MDCK cells was altered by subjecting cells to variants of Ringer’s solution as described previously^44^. Cells were incubated with either acetate Ringer’s solution (80 mM NaCl, 70 mM sodium acetate, 5 mM KCl, 2 mM CaCl_2_, 1 mM MgCl_2_, 2 mM NaH_2_PO_4_, 10 mM HEPES, and 10 mM glucose, final pH 6.9) or Alkaline Ringer’s solution (155 mM NaCl, 5 mM KCl, 2 mM CaCl_2,_ 1 mM MgCl_2_, 2 mM NaH_2_PO_4,_ 10 mM HEPES, 10 mM Glucose, 0.5 mg mL^-1^ BSA and 20 mM NH_4_Cl) for 30 minutes before lifting the culture insert. After lifting the confinement, the cells were allowed to migrate in either Acetate Ringer’s or Alkaline Ringer’s and migration was followed up to 4 hours. Control cells migrated in the presence of Opti-MEM, which has a pH of 7.2, and its composition overlaps with that of Ringer’s solution. Cells were live imaged using 20X objective on Leica DMI8 inverted microscope using Leica Application Suite X (LAX, v3.7.0.20979) under different treatment conditions.

### Antibodies and Plasmids

Source and dilution information for all primary and secondary antibodies used in immunofluorescence staining are given in Supplementary Table 1. Details of plasmids used in this study are listed in Supplementary Table 2 with their source.

### Micropatterning

PDMS stencils with pillars of defined shapes were created using soft photolithography. Briefly, 150 um thick SU-8 2075 (Kayaku advanced materials, Y111074) was coated on a silicon wafer followed by soft baking at 65° C for 5 mins and 95° C for 20 mins. SU-8 was exposed in the desired pattern using the MicroWriter ML 3 Pro (Durham Optics Magneto Ltd) followed by post-exposure baking at 65° C for 7 mins and 95° C for 15 mins. The unexposed photoresist was removed by immersing the wafers in the SU-8 developer (Kayaku advanced materials, Y020100) for 8-10 mins. Patterned SU-8 wafers were then hard-baked at 170° C for 45 mins. PDMS (Sylgard 184, Dow Corning) was mixed in a ratio of 1:10, degassed, and poured over the patterned wafers. After curing at 85 °C for 2 h, stencils with pillars were peeled off. The stencils were then treated with 2% Pluronic acid for 2 hrs followed by 15 mins of 1X PBS wash. PDMS stencil and a petri dish were plasma cleaned for 10 seconds and the stencil was then placed upon the dish. Cells were seeded around the pillars, and after confluence, the stencils were removed allowing the cells to migrate into the gaps.

### Image Analysis

All image analysis for this study was performed using Fiji. Peripheral occupancy Fraction (POF) was measured for cells at the margin of a migrating collective. The ROI box with a fixed width of 2.5 μM from the cell edge facing the gap was drawn over the cell of interest. The box was duplicated for the lysosome channel and a threshold was established to mark lysosomes. Thresholding for lysosomes was done using the Moments function of Fiji and it was kept constant across experiments. Using the Analyze Particles tool of Fiji, the lysosome count was determined. For each cell, the total number of lysosomes was also calculated by manually marking the cell of interest and following the same approach as mentioned above for lysosome number calculation in the ROI. For calculating the POF, the number of lysosomes in ROI was divided by the total number of lysosomes in that cell. 20 marginal and ∼10 leader cells per experiment were quantified from three independent experiments. For calculating the junctional occupancy fraction for lysosomes (Fig 1E), we measured the number of lysosomes in the cell-cell junctions of the leader cells and non-leader marginal cells by drawing an ROI of 2.5 μM from the junction, followed by dividing it to the total number of lysosomes in the cell. For measuring the POF for different organelles, we followed the same approach, but instead of the count of the organelle, we used the intensity in ROI and the total intensity of the organelle of interest.

To measure POF for experiments involving OptoGEF-RhoA, cells adjoining the photoactivated cell were marked manually using the DIC channel to mark cell boundaries. The image was duplicated for the lysosome channel. Lysosome intensity was determined using Fiji. This was followed by marking the cell periphery of the same cell by using the Edit → Selection → Enlarge tool of Fiji to reduce the outline by 2.5 μm. Using the Edit → Clear tool the lysosomes in the perinuclear region were omitted and the intensity of remaining lysosomes in the cell periphery was determined. To get the POF for cells, the intensity of lysosomes was divided by the total cell intensity of lysosomes for each cell. POF was determined for pre-activation and post-activation states. Change in POF was compared between pre-and post-activation for the same cell and 25-27 cells were analysed per treatment from three independent experiments.

Colocalization analysis of different forms of small GTPase Rac1 with lysosomal marker LAMP1 was performed using the JACoP tool of Fiji. For measuring Mander’s coefficient, cytoplasmic expression was analysed by marking the inside of the cell boundary using a freehand tool in Fiji. The cytoplasmic Rac1 expression includes its vesicular component and its co-localization with lysosomes was measured. Analysis was performed for three independent experiments and 20 cells per transfection per experiment were analysed.

Tracking of cells during collective migration was done using the Manual Tracking tool of Fiji. For Fig. 4, the net displacement of was calculated from the results obtained by tracking cells that emerge as leader cells from T-0 to T-2hrs post confinement lift-off. We tracked 20 leader cells over 3 independent experiments. We followed the same manual tracking approach for tracking cells with biased lysosome positioning (Cell lines expressing mch-Kif5b*-Strep, Strep-KifC1*-mCh co-transfected with LAMP1-SBP-GFP plasmids). To measure lysosome speed from timelapse videos, TrackMate tool of Fiji was used with the following parameters: Vesicle diameter-1μm; Detector-DoG; Initial thresholding-none; Tracker-Simple LAP tracker; Linking max distance-2 μm; Gap-closing max distance-2 μm, Gap-closing max frame gap-2; Filters-none.

### Statistical analysis

Statistical analyses were carried out in GraphPad Prism 9. Statistical significance was calculated by Unpaired t-test with Welch’s correction or One-way ANOVA as mentioned in the corresponding figure. Scatter-bar plots were displayed as mean ± s.e.m. p-values greater than 0.05 were considered to be statistically not significant. No statistical methods were used to set the sample size. Quantification was done using data from at least three independent biological replicates. For analysis involving live-imaging experiments, data were collected from three independent experiments. All the experiments with representative images were repeated at least thrice.

### AUTHOR CONTRIBUTIONS

R.M. and T.D. formulated the project. R.M. performed the majority of the experiments and analysis. S.R. exclusively performed and analysed micropatterning experiments and contributed to other experiments as well. P.K. performed opto-GEF experiments. S.B. and D.M. performed mouse embryonic skin wound assays. M.J. provided critical inputs to the design of *Drosophila* embryo experiments. R.M. and T.D. wrote the manuscript. M.J. edited the manuscript. All authors agreed on the manuscript as in the submitted version.

## Supporting information

Combined Supplementary Information

## ACKNOWLEDGEMENT

We thank Ullas Kolthur and Aneesh T. Veetil for the critical discussion. R.M. is grateful to Praver Gupta for his critical inputs on POF analysis and image representation. T.D. is a DBT/Wellcome Trust India Alliance intermediate fellow and partner group leader of the Max Planck Society (MPG), Germany. R.M. is a DBT/Wellcome Trust India Alliance early career fellow. Authors sincerely acknowledge generous funding from the DBT/Wellcome Trust India Alliance (Ref. No. IA/E/19/1/504967 to R.M. and IA/I/17/1/503095 to T.D.), partner group grant of the Max Planck Society, and intramural funds at TIFR Hyderabad from the Department of Atomic Energy (DAE), India, under the Project Identification Number RTI 4007, towards supporting this project.

