## Supplementary material for "Mechanosensitive dynamics of lysosomes along microtubules regulate leader cell emergence in collective cell migration": Combined Supplementary Information

Marwaha *et al*.

This document contains:

1. Supplementary Table 1: Antibody information
2. Supplementary Table 2: Plasmid information
3. Supplementary Figure Legends
4. Supplementary Video Legends
5. Supplementary Figures 1-8

**1. Supplementary Table 1: Antibody information**

| *S.no.* | *Antibody* | *Cat.no.* | *Dilution* |
|---|---|---|---|
| 1. | Anti-LAMP1 | ab24170 (Abcam) | 1:2500 |
| 2. | Anti-LAMP2 | MCA2558GA (BioRad) | 1:500 |
| 3. | Anti-LAMP1 | ab30687 (Abacm) | 1:200 |
| 4. | Anti-Paxillin | ab32084 (Abcam) | 1:200 |
| 5. | Anti-TFEB | 37785S (CST) | 1:200 |
| 6. | Anti-LAMTOR4 | 13140S (CST) | 1:200 |
| 7. | Anti-Tom20 | 42406S (CST) | 1:200 |
| 8. | Anti-Rab7 | 9367T (CST) | 1:200 |
| 9. | Anti-EEA1 | 610456 (BD Biosciences) | 1:250 |
| 10. | Alexa fluor conjugated secondary antibodies | CST | 1:500 |
| 11. | Alexa fluor Phalloidin | CST | 1:1000 |

**2. Supplementary Table 2: Plasmid information**

| *S.no.* | *Plasmid* | *Source* |
|---|---|---|
| 1. | mCherry-Kif5b*-Strep | Addgene |
| 2. | Strep-KifC1*-mCherry | Addgene |
| 3. | LAMP1-SBP-GFP | Addgene |
| 4. | ARHGEF11(DHPH)-CRY2- mCherry (optoGEF-RhoA) | Gift from Dr. Xavier Trepat |
| 5. | mito-CIBN-GFP | Gift from Dr. Xavier Trepat |
| 6. | CIBN-CAAX | Addgene |
| 7. | pcDNA3-EGFP-Rac1 WT | Addgene |

| | | |
|---|---|---|
| 8. | pcDNA3-EGFP-Rac1 CA (Q61L) | Addgene |
| 9. | pcDNA3-EGFP-Rac1 DN (T17N) | Addgene |
| 10. | mEos-CA Rac1 | Addgene |
| 11. | mEos-DN Rac1 | Addgene |
| 12. | GFP-KLC | Gift from Dr. Mahak Sharma |
| 13. | FLAG-SKIP WT | Gift from Dr. Mahak Sharma |
| 14. | FLAG-SKIP WD->2XA | Gift from Dr. Mahak Sharma |
| 15. | pcDNA3-EGFP-RhoA CA (Q63L) Addger | Addgene |
| 16. | pcDNA3-EGFP-RhoA DN (T19N) | Addgene |
| 17. | pECFP-Cdc42 CA (Q61L) | Addgene |
| 18. | pcDNA3-EGFP-Cdc42DN (T17N) | Addgene |

### 3. SUPPLEMENTARY FIGURE LEGENDS

**Supplementary Figure 1: Lysosome distribution in marginal cells during wound closure in different model systems. A**) *Drosophila melanogaster* stage 15 embryos were wounded using glass needles, PFA fixed 15mins post wounding and immunostained with anti-LAMP1 to mark lysosomes, Alexa-fluorophore conjugated phalloidin to mark actin and DAPI to stain the nucleus. **B**) Representative confocal micrographs and 3-dimensional volume view of mouse embryonic skin wound allowed to heal for 30mins post incision, fixed and immunostained for lysosomes (Anti-LAMTOR4) and actin (Alexa fluorophore-conjugated phalloidin). **C**) Schematic representation of 2-dimensional gap closure assay used in this study to look at collective migration of mammalian epithelial cells. **D**) Representative confocal micrographs of MDCK cells allowed to migrate for T-120mins post confinement lift-off, PFA fixed and immunostained for actin (Alexa fluorophore-conjugated phalloidin) and cortactin. Yellow arrowheads mark a leader cell having lamellipodia and enlarged cell size. Scale bar, 10 µm for all sub-figures; ** corresponds to $p<0.01$ and *ns* is not significant.

**Supplementary Figure 2: Lysosome accumulation in cells at the wound margin during collective migration. A**) Representative confocal images of collectively migrating MDCK cells for indicated time points. Lysosomes accumulate in the direction of migration in cells at the wound margin. **B**) Fraction of peripheral lysosomal accumulation in cells at the wound migration for T-0,15,30,60,120, and 240mins was measured by calculating the POF of lysosomes and represented here as a column graph (n=3, 15 cells per experiment were analysed). Data are mean±s.e.m. **C**) Heat map representing the correlation between POF of lysosomes and displacement of MDCK cells migrating collectively at T-0, 15, 30, 60, 120 and

240mins post confinement lift-off. The scale bar represents normalized percentage values of POF of lysosomes and migration distance. Values at 240 mins were considered as 100% and values at other time points were normalized to this. **D**) Representative confocal micrographs of EPH4-Ev cells undergoing collective migration. The cells were PFA fixed at T-0 and T-120mins post confinement lift-off, immunostained for lysosomes (Anti-LAMP1), actin (Alexa fluorophore-conjugated phalloidin) and nucleus (DAPI). **E**) POF of lysosomes was calculated for marginal and leader cells during collective migration of EPH4-Ev cells (n=3, 15 cells for each type and ~9 leader cells per experiment were analysed. Data are mean±sem. Leader cells have a higher POF of lysosomes as compared to non-leader marginal cells. Statistical significance was calculated using Student's t-test with Welch's correction. Scale bar, 10 µm for all sub-figures. *, ** signifies p-value<0.05 and <0.01 respectively, *ns* is not significant.

**Supplementary Figure 3: Enhanced lysosome accumulation to cell periphery is driven by organelle movement on microtubule tracks. A**) TFEB localization in a migrating monolayer of MDCK cells. MDCK cells were fixed and immunostained for total TFEB, actin, and nucleus post 120mins of migration, and images were acquired using a confocal microscope. **B**) Column graph of the total number of lysosomes in marginal (both non-leaders and leaders) and follower cells measured using Fiji's Analyze particle tool (n=3; 20 cells per type per experiment were analysed. Data are mean±sem). **C**) Microtubule and lysosome localization in collectively migrating EPH4-Ev cells. EPH4-Ev cells seeded in migration chambers were allowed to migrate post-confinement lift-off for the indicated time points, PFA fixed and immunostained with anti-LAMP1, anti-α-tubulin, Alexa-fluorophore conjugated phalloidin to mark actin and DAPI to stain the nucleus. Images were acquired on a confocal microscope. **D**) MDCK cells migrated for the indicated time points, PFA fixed and immunostained for lysosomes (Anti-LAMP1) and microtubules (Anti-Alpha Tubulin). Accumulation of microtubules and lysosomes at the cell periphery can be observed and simultaneous at the indicated time points. Scale bar, 10 µm for all sub-figures. **** signifies p-value<0.0001 and *ns* is not significant.

**Supplementary Figure 4: Lysosome accumulation to cell periphery is specific as other cellular organelles do not polarize during collective cell migration. A-E**) Representative confocal micrographs of MDCK cells at T-0 and T-120mins of migration immunostained with indicated antibodies marking lysosomes (**A**), late-endosomes (**B**), early endosomes (**C**), stable cell line expressing mcherry-Sec61 to mark endoplasmic reticulum (**D**) and mitochondria (**E**). Insets show leader-like/leader cells emerging from a collectively migrating epithelia. **F**) Scatter column graph showing POF of the indicated organelles, calculated by taking a ratio of organelle marker fluorescence intensity in ROI to the total intensity in the cell (n=3, 30 cells per experiment for each organelle). Student t-test with Welch's correction was used for statistical analysis. Scale bar, 10 µm for all sub-figures, *** signify p-value<0.001 and *ns is not significant.*

**Supplementary Figure 5: Lysosome positioning dictates leader cell emergence. A**) Representative confocal micrographs of control (optiMEM), AR and AlkR treated MDCK cells allowed to migrate for T-120 mins post confinement lift-off. Lysosome distribution was observed by immunostaining for anti-LAMP1 and actin was labelled by Alexa-fluorophore conjugated phalloidin. **B-D**) Representative time stamps of Control, Acetate Ringer's (AR), and Alkaline Ringer's (AlkR) solution-treated MDCK cells. Cells migrated for indicated time points and were observed for leader cells' emergence, identified by the extended lamellipodial structure under different conditions. Yellow arrowheads mark emerging leader cells. **E**) Scatter plot representing the fraction of cells under Control, AR, and AlkR treatment emerging as leader cells, identified by the presence of extended lamellipodia (n=3; 12 wound margin regions comprising of an average 70 cells per margin were analysed for each treatment per experiment). **F-G**) Representative confocal micrographs of MDCK stable cell lines expressing mCh-Kif5b*-Strep (**F**) and Strep-Kifc1*-mCh (mCh: mCherry) (**G**), respectively. **H-I**) EPH4-Ev cells transiently co-transfected with LAMP1-SBP-GFP and either mCh-Kif5b*-Strep (**H**) or Strep-Kifc1*-mCh (**I**) respectively and allowed migration for 240 mins. The presence or absence of lamellipodia and leader cell emergence in co-expressing cells was assessed by confocal microscopy. Scale bar, 10 μm for all sub-figures; * and **** signify p-value<0.05 and 0.0001 respectively.

**Supplementary Figure 6: Actomyosin contractility inhibition leads to changes in lysosome spatial dynamics. A-C**) Time stamps of control (**A**), blebbistatin (**B**), and Y27632 (**C**) treated MDCK cells labelled for lysosomes using Dextran-OregonGreen and SiR-Actin. Cells were allowed to migrate for 120 mins and live-imaged using a confocal microscope. White arrowheads in (A) mark peripheral lysosomes in an emerging leader cell. Cyan arrowheads in (B) and (C) mark tubular lysosomes emerging upon actomyosin contractility inhibition. **D**) Scatter plot showing the speed of lysosomes (μm/sec) in control, blebbistatin and Y27632 treated cells. Scale bar-10μm. **** represents p-value<0.0001 and *ns* is not significant.

**Supplementary Figure 7: Lysosome distribution depends upon cellular forces generated by the actomyosin cytoskeleton. A**) Schematic representation of the forces acting on a pair of cells. **B**) MDCK cells were seeded at confluency to form cell pairs. Cells were incubated with DMSO (Control) or Blebbistatin overnight, followed by blebbistatin washout for 240 mins for one set. Lysosome distribution was analysed using confocal microscopy post-fixation and immunostaining with anti-LAMP1 and anti-LAMP2. **C**) Diagram representing Opto-RhoAGEF MITO-CIBN-CAAX expression in cells and its localization upon photoactivation by 488nm light. Opto-RhoAGEF MITO-CIBN-CAAX upon activation localizes to mitochondria, thus causing no change in cell contractility. **D**) Live-imaging snapshots of MDCK cells expressing Opto-RhoAGEF MITO-CIBN-CAAX before and after photoactivation. Montage of inverted lysosome signal in 2.5μm wide cell periphery region. No significant increase in lysosome signal is observed post-activation. **E**) Change in POF of lysosomes was calculated for cells juxta-positioned to optogenetically altered cells and is represented here in this column graph (n=3, 27 cells were analysed). Statistical analysis was performed using the Student's t-test with Welch's correction. **F**) Brightfield images showing MDCK cells seeded in single-beak micropatterned chambers. **F**) Live imaging snapshots of LifeAct-MDCK cells seeded in single-beak micropatterned chambers and allowed to migrate post-confinement lift-off. The cells close the wound gap. Scale bar, 10 μm for all sub-figures, *ns* is not significant.

**Supplementary Figure 8: Screening of Rho family GTPases. A-D**) MDCK cells were transiently transfected with the indicated plasmids and allowed to migrate for 240 mins. Post-migration cells were fixed and stained for lysosomes (anti-LAMP1), and actin (Phalloidin). Confocal imaging was performed to analyse the co-localization of lysosomes with GFP- RhoA CA, GFP-RhoA DN, CFP-CDC42 CA and CFP-CDC42 DN. **E-G**) Representative confocal micrographs of EPH4-Ev cells transiently expressing GFP-Rac1 WT (**E**), GFP-Rac1 CA (**F**) and GFP-Rac1 DN (**G**), allowed to migrate for 240 mins, fixed and immunostained for lysosomes (anti-LAMP1) for co-localization studies. **H**) Co-localization of Rac1 WT, DN and CA with LAMP1 was analysed by measuring Mander's coefficient, represented here as a scatter column graph (n=2; 10 cells per experiment). Data are mean±s.d.

### 4. SUPPLEMENTARY VIDEO LEGENDS

**Supplementary video 1:** Lysosome and actin dynamics in migrating LifeAct-GFP MDCK cells. Dextran-647-labelled lysosomes (magenta) in emerging leaders marked by pink arrowhead show higher bidirectional movement as compared to lysosomes in neighbouring marginal cell marked by yellow arrowhead. The movie is shown at seven frames per second.

**Supplementary video 2:** Growth of actin structure depends upon lysosome enrichment within a leader cell. LifeAct-MDCK cells (green) and lysosomes (magenta) were live imaged under a confocal microscope. The movie is shown at ten frames per second.

**Supplementary video 3:** Peripheral lysosome bias enhances leader cell emergence. MDCK cells stably expressing mCherry-Kif5b*-Strep transiently transfected with LAMP1-SBP-GFP were imaged using fluorescence microscope. Blue and green line marks the track of an emerging leader cell. The movie is shown at seven frames per second.

**Supplementary video 4:** Cells with perinuclear lysosome accumulation do not emerge as leaders. MDCK cells stably expressing Strep-Kifc1*-mCh transiently co-expressed with LAMP1-SBP-GFP were imaged using fluorescence microscope. Green dot and line mark the track of co-transfected cell. The movie is shown at seven frames per second.

**Supplementary video 5:** Lysosome dynamics upon actomyosin contractility inhibition. Control, Blebbistatin and Y27632 treated MDCK cells were allowed to migrate for 120mins post confinement lift-off. SiR-Actin and Dextran-Oregon Green-labelled lysosome dynamics were recorded live on a confocal microscope. Cells were imaged for 5 mins at an interval of 5 secs. The movie is shown at 10 frames per second.

**Supplementary video 6:** Time-lapse video of MDCK cells expressing Opto-CIBN-GEF-RhoA. The cells were imaged live on a confocal microscope both pre-activation and post-activation for 5mins with 5secs interval. Video is shown at seven frames per second.

**Supplementary video 7:** Time-lapse video of MDCK cells expressing Opto-GEF-RhoA MITO-CIBN. The cells were imaged live on a confocal microscope both pre-activation and post-activation for 5mins with 5secs interval. Video is shown at seven frames per second.

**Supplementary video 8:** Gap closure of the micropatterned wound. Post confinement lift-off LifeAct-GFP MDCK cells seeded in a single beak micropattern chamber migrate and close the wound. The movie is shown at 10 frames per second.

**Supplementary video 9:** Collective migration of LifeAct-GFP MDCK cells using single-beak micropatterned chamber. Dynamics of actin (green) and dextran-647-labelled lysosomes (magenta) were recorded at an interval of 3 mins 25 secs. The movie is shown at seven frames per second.

**Supplementary video 10:** Localization dynamics of inactive Rac1. MDCK cells transiently transfected with mEos-Rac1-DN and dextran-647 labelled lysosomes were subjected to photoconversion and further imaged live for 5mins to observe the co-localization between inactive Rac1 and lysosomes in migrating cells. The movie is shown at seven frames per second.

**Supplementary video 11:** Localization dynamics of active Rac1. MDCK cells transiently transfected with mEos-Rac1-CA and dextran-647-labelled lysosomes were subjected to photoconversion and further imaged live for 5 mins to observe the dynamics of active Rac1 and lysosomes in migrating cells. The movie is shown at seven frames per second.

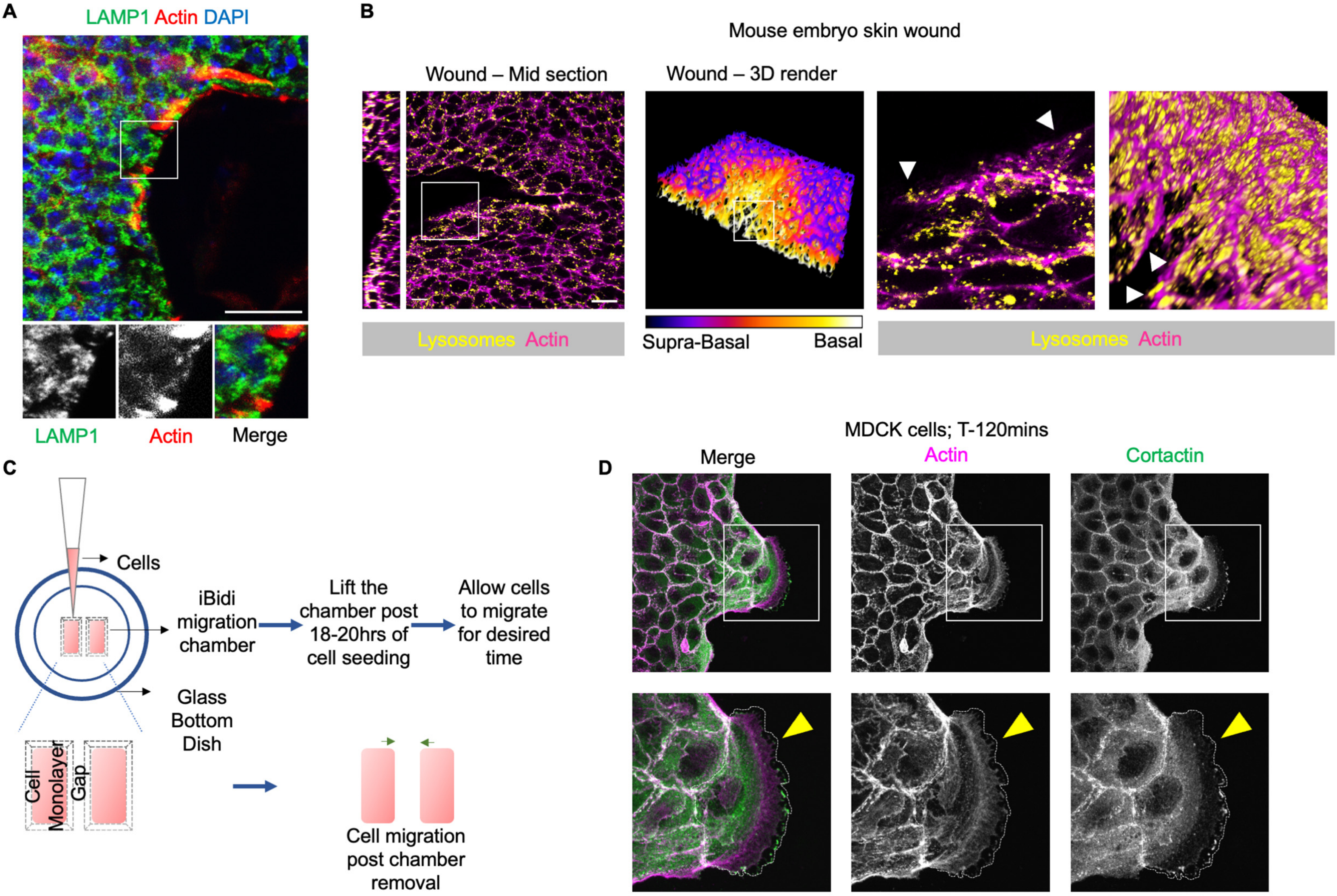


**Supplementary Figure 1**

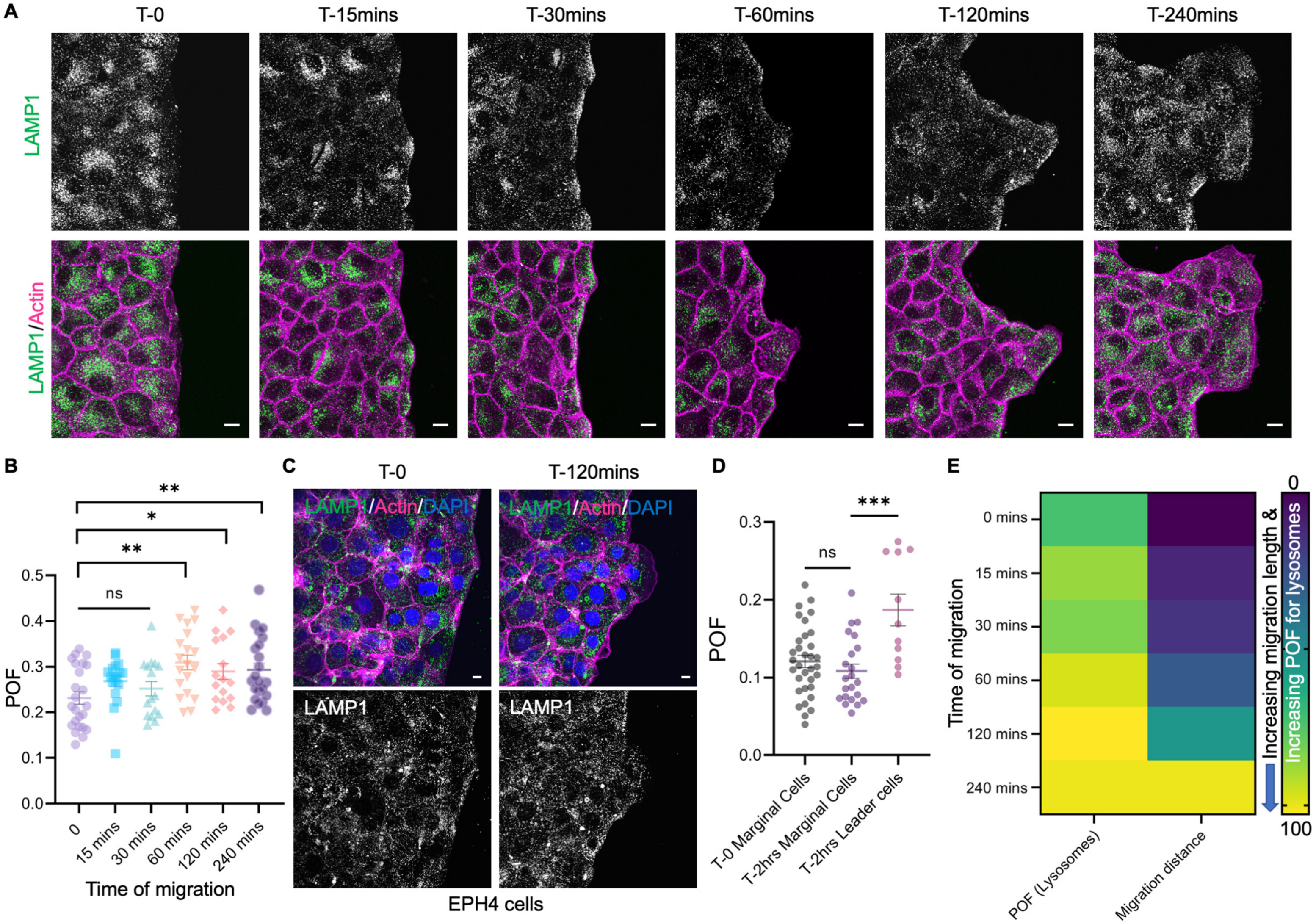


**Supplementary Figure 2**

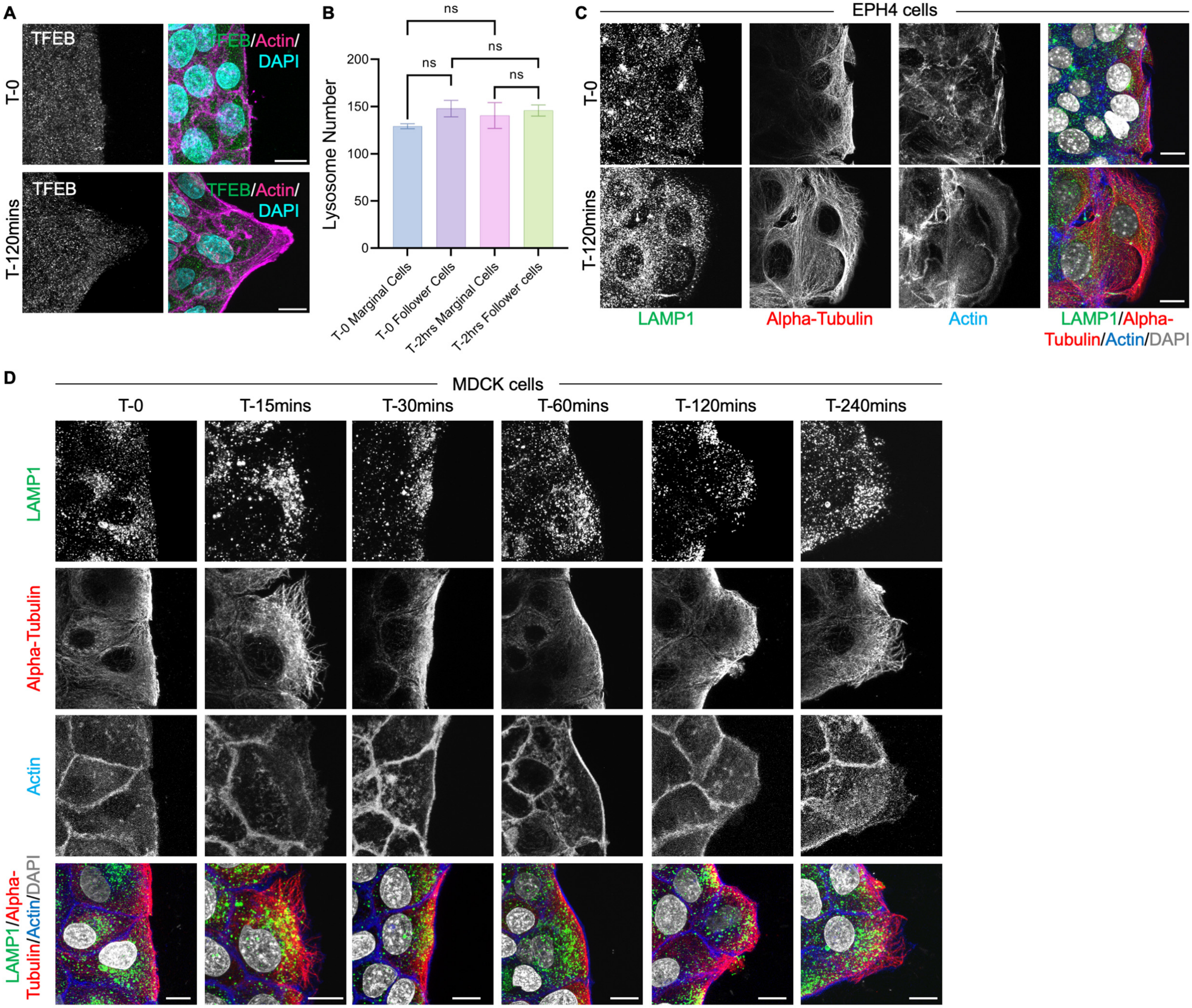


**Supplementary Figure 3**

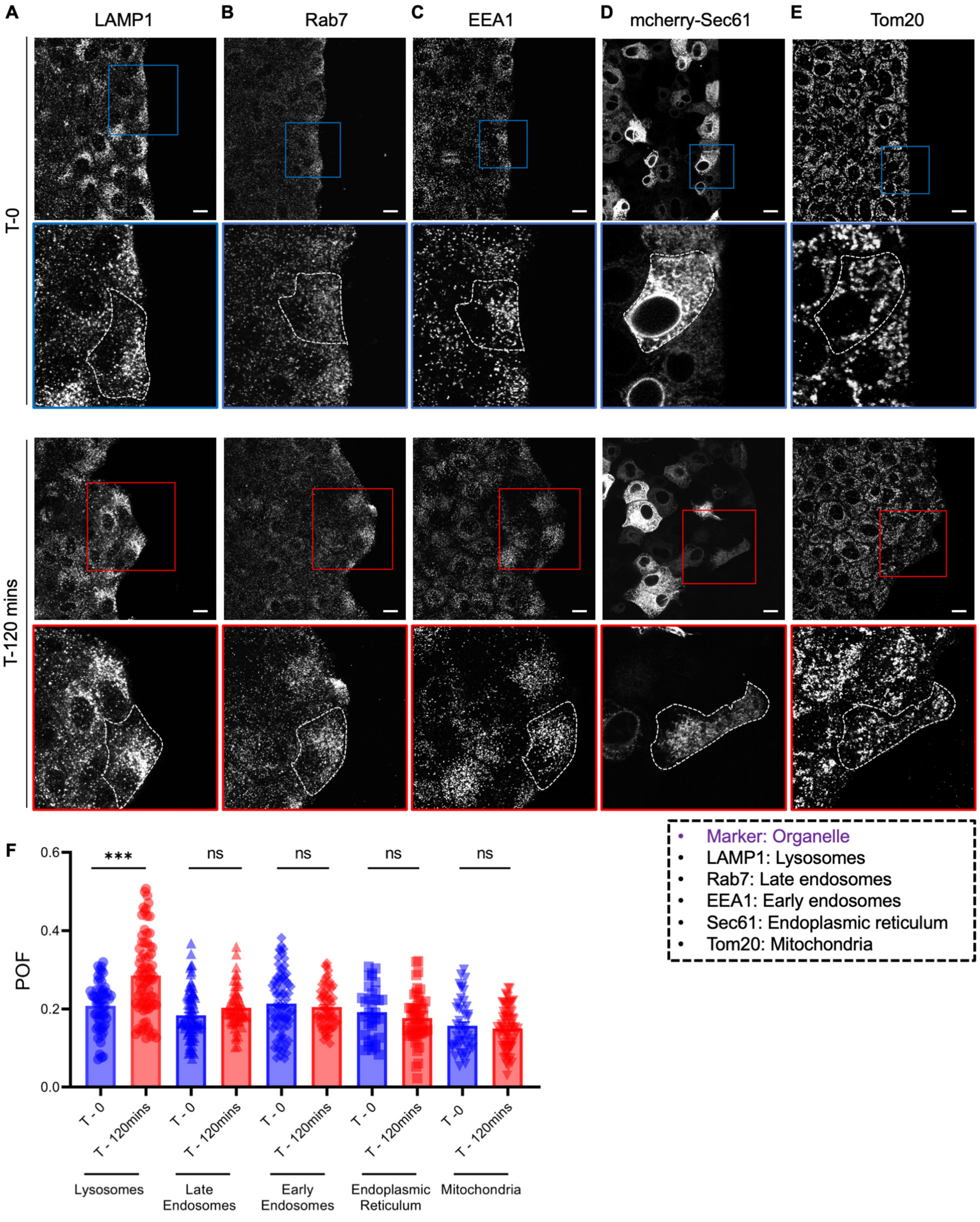


**Supplementary Figure 4**

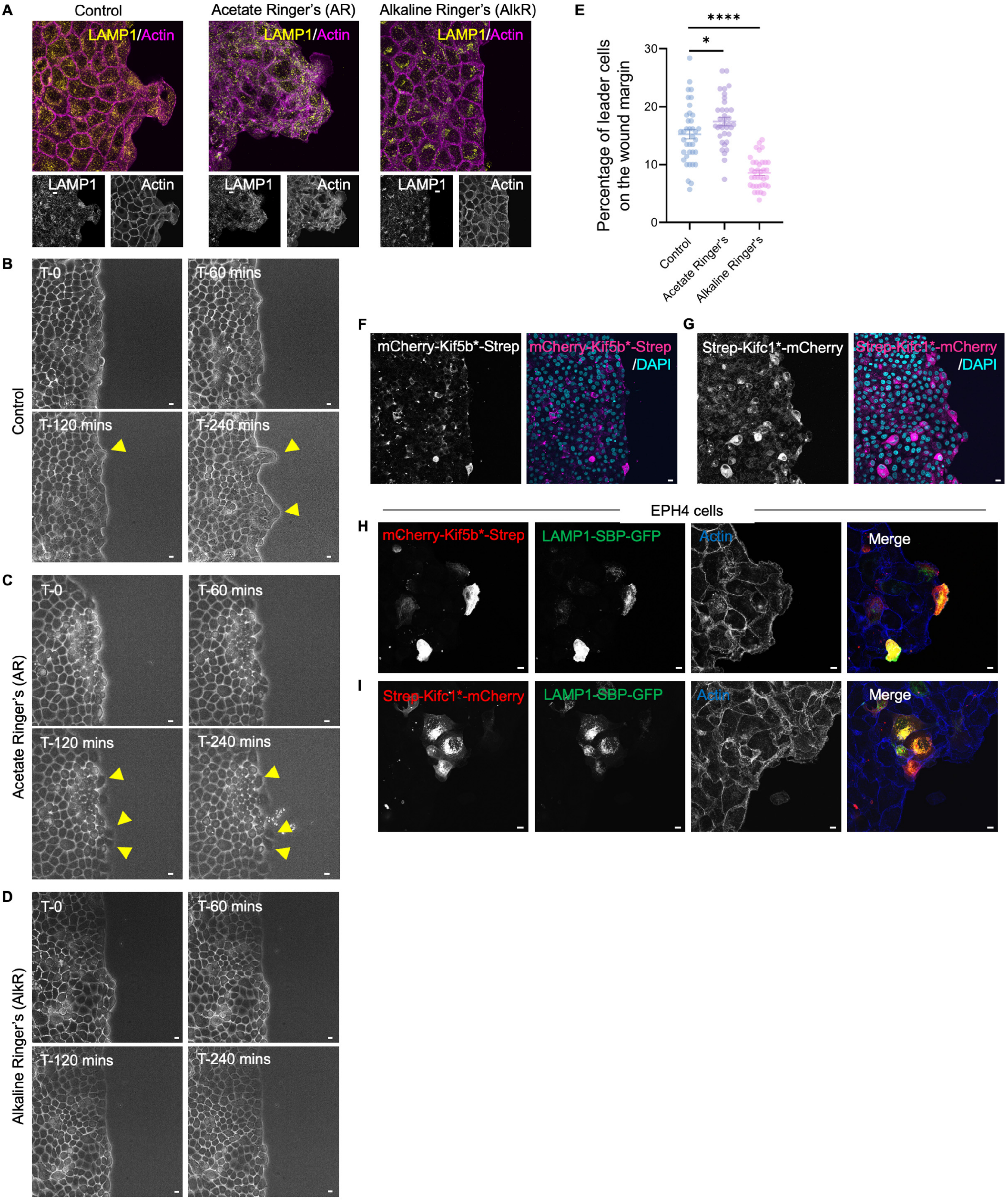


**Supplementary Figure 5**

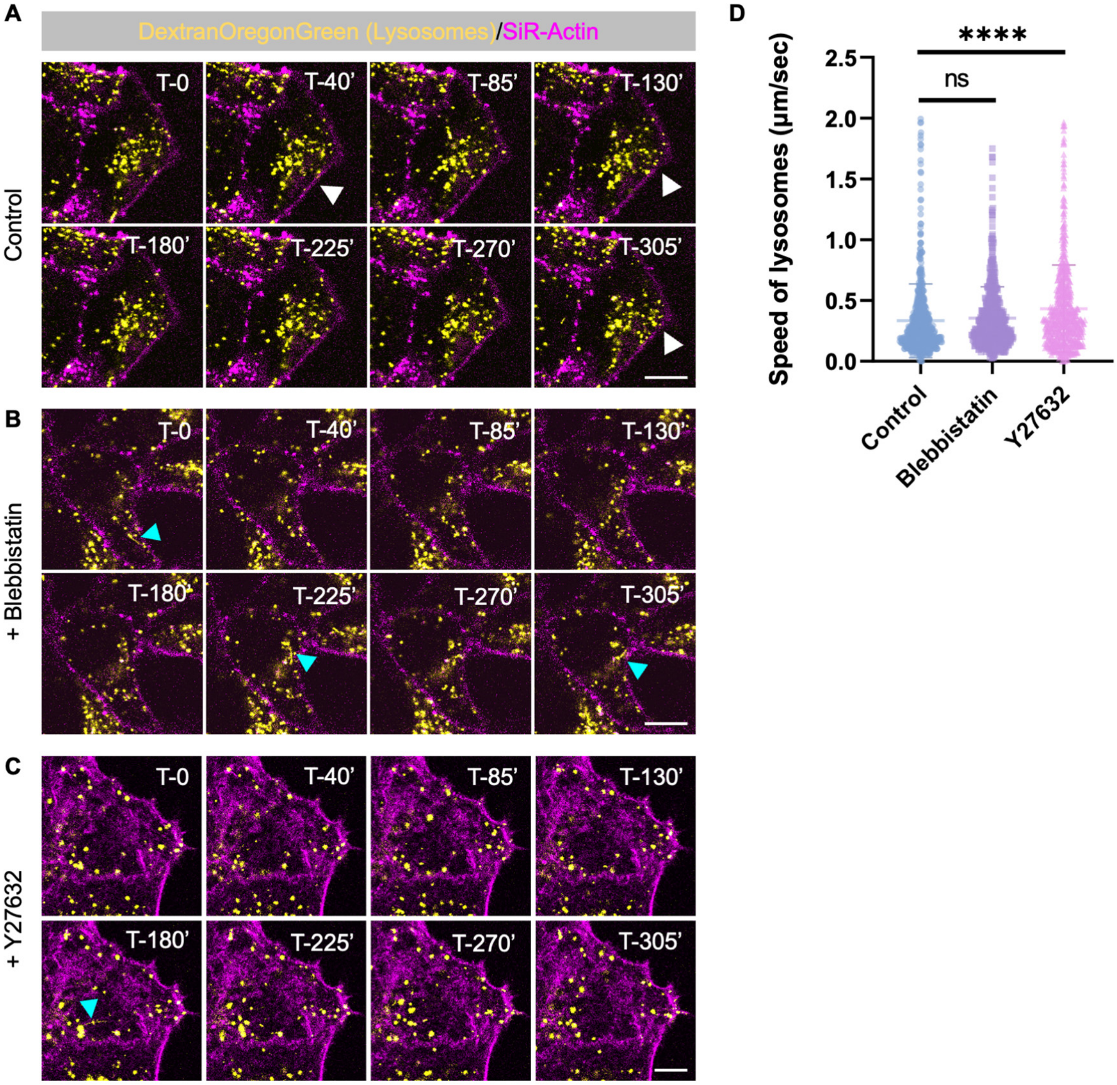


Supplementary Figure 6

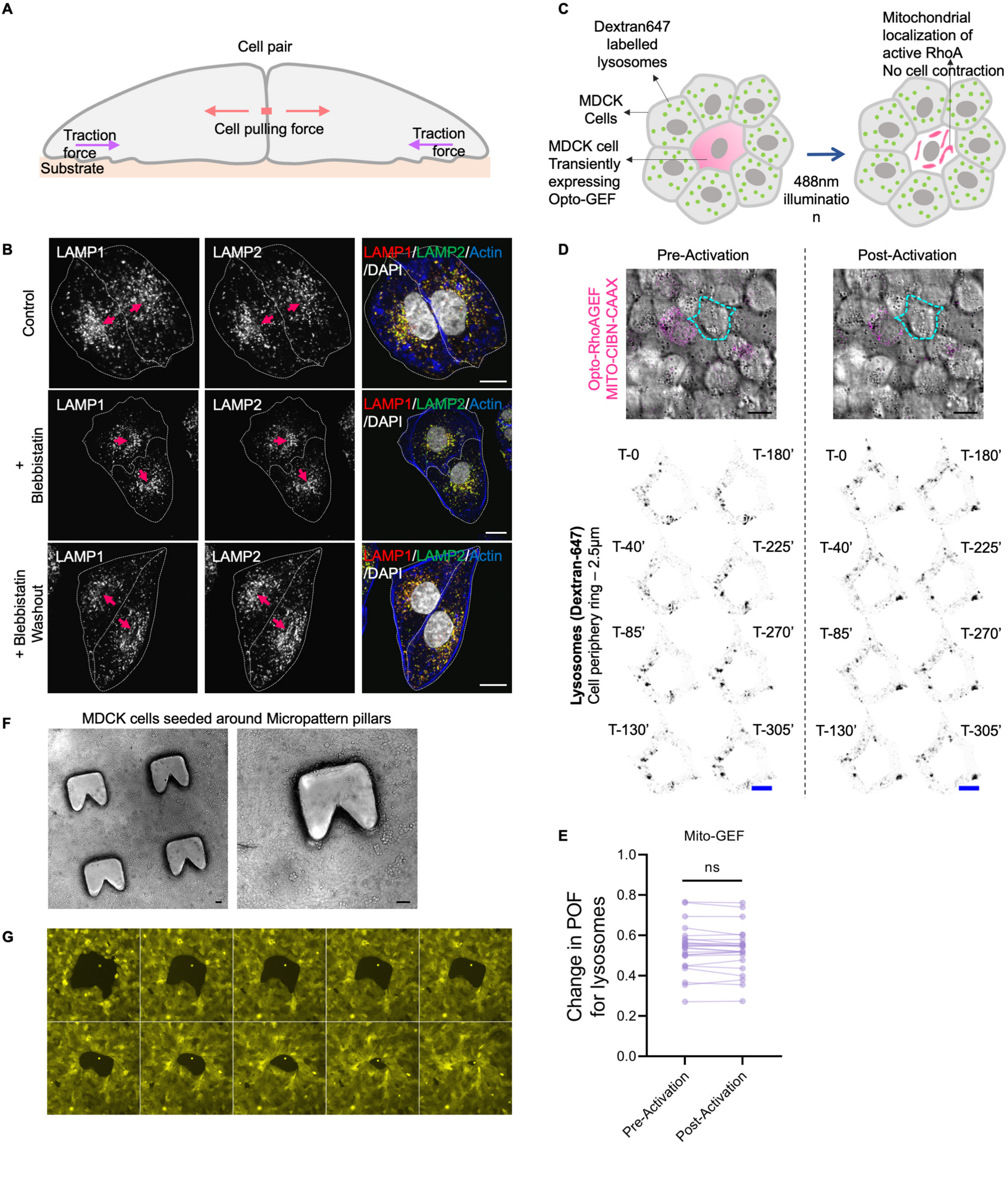


**Supplementary Figure 7**

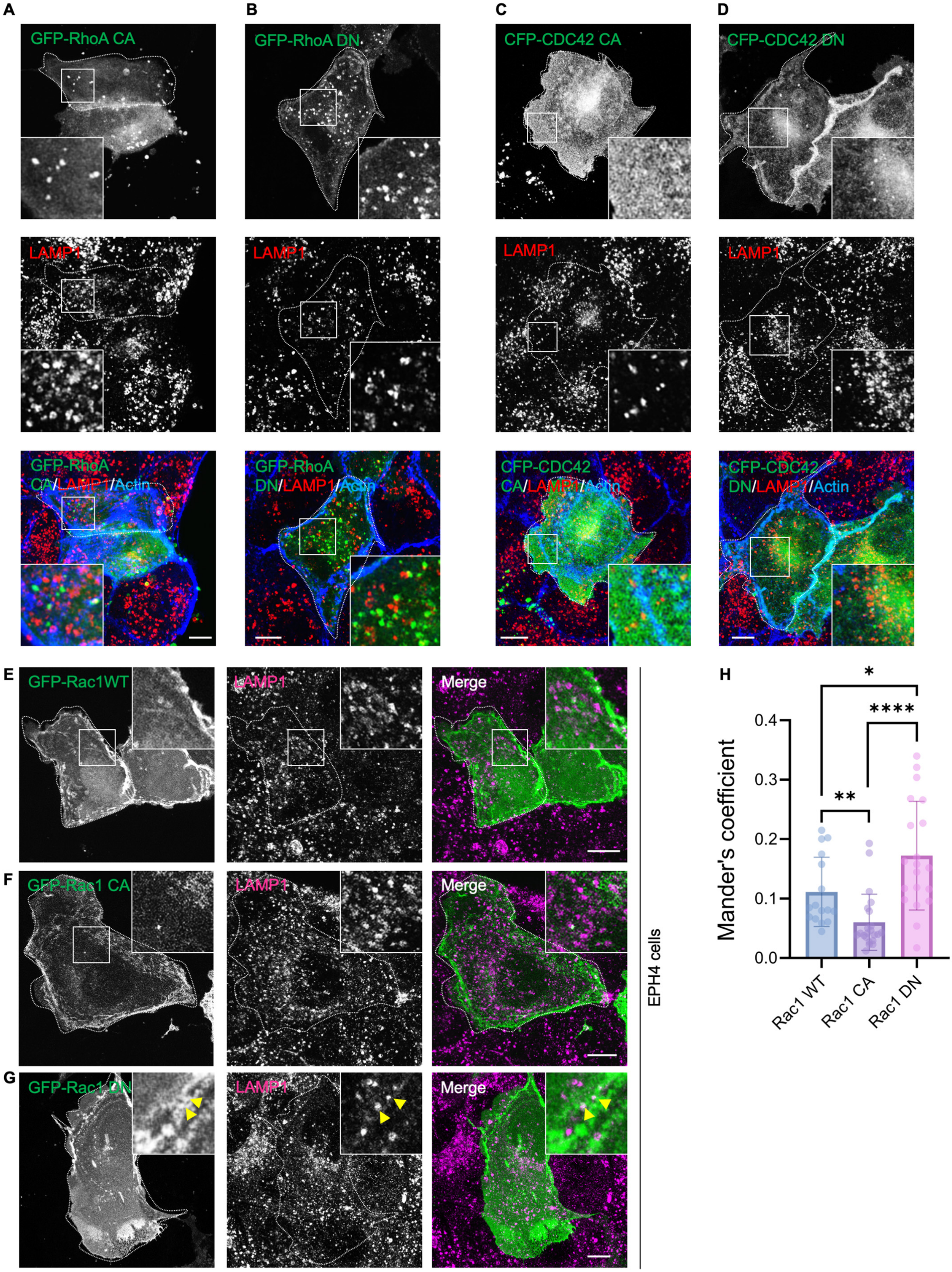


**Supplementary Figure 8**
